# MCC950/CRID3 potently targets the NACHT domain of wildtype NLRP3 but not disease-associated mutants for inflammasome inhibition

**DOI:** 10.1101/634493

**Authors:** Lieselotte Vande Walle, Irma B. Stowe, Pavel Šácha, Bettina L. Lee, Dieter Demon, Amelie Fossoul, Filip Van Hauwermeiren, Pedro H. V. Saavedra, Petr Šimon, Vladimír Šubrt, Libor Kostka, Craig E. Stivala, Victoria C. Pham, Steven T. Staben, Sayumi Yamazoe, Jan Konvalinka, Nobuhiko Kayagaki, Mohamed Lamkanfi

## Abstract

The NLRP3 inflammasome drives pathological inflammation in a suite of autoimmune, metabolic, malignant and neurodegenerative diseases. Additionally, *NLRP3* gain-of-function point mutations cause systemic periodic fever syndromes that are collectively known as cryopyrin-associated periodic syndromes (CAPS). There is significant interest in the discovery and development of diarylsulfonylurea Cytokine Release Inhibitory Drugs (CRIDs) such as MCC950/CRID3, a potent and selective inhibitor of the NLRP3 inflammasome, for the treatment of CAPS and other diseases. However, drug discovery efforts have been constrained by the lack of insight in the molecular target and mechanism by which these CRIDs inhibit the NLRP3 inflammasome. Here, we show that the NACHT domain of NLRP3 is the molecular target of diarylsulfonylurea inhibitors. Interestingly, we find photoaffinity labelling of the NACHT domain requires an intact (d)ATP-binding pocket and is substantially reduced for most CAPS-associated NLRP3 mutants. In concordance, MCC950/CRID3 failed to inhibit NLRP3- driven inflammatory pathology in two mouse models of CAPS. Moreover, it abolished circulating levels of interleukin (IL)-1β and IL-18 in LPS-challenged wildtype mice but not in *Nlrp3*^L351P^ knock-in mice and ex vivo-stimulated mutant macrophages. These results identify wildtype NLRP3 as the molecular target of MCC950/CRID3, and show that CAPS-related NLRP3 mutants escape efficient MCC950/CRID3 inhibition. Collectively, this work suggests that MCC950/CRID3-based therapies may effectively treat inflammation driven by wildtype NLRP3, but not CAPS-associated mutants.

## Introduction

Inflammasomes are a suite of multi-protein complexes that play central roles in innate immune responses through their ability to recruit and activate caspase-1 (Broz and Dixit, 2016; Lamkanfi and Dixit, 2014). This cysteine protease cleaves the cytokines interleukin (IL)-1β and IL-18 and drives pyroptosis, a highly inflammatory regulated cell death mode that is induced by cleavage of gasdermin D (GSDMD) (Kayagaki et al., 2015; Shi et al., 2015). Amongst the different inflammasome pathways, the NLRP3 inflammasome responds to the broadest suite of inflammasome agonists that includes diverse pathogen-associated molecular patterns (PAMPs), host-derived danger-associated molecular patterns (DAMPs) like ATP, protein aggregates and β-fibrils such as β-amyloid, a broad range of environmental insults and ionophores such as nigericin and medically relevant crystals such as alum, CCPD, MSU, silica and asbestos (Broz and Dixit, 2016; Lamkanfi and Dixit, 2014). Moreover, the NLRP3 inflammasome is engaged by Gram-negative pathogens, lipopolysaccharides (LPS) of which are sensed in the cytosol by the non-canonical NLRP3 inflammasome pathway (Hagar et al., 2013; Kayagaki et al., 2011; Kayagaki et al., 2013; Shi et al., 2014). Through the latter mechanism, cleavage of GSDMD by caspase-11 – and its human orthologs caspases 4 and 5 – promotes assembly of cytolytic GSDMD pores in the plasma membrane that also activate the NLRP3 inflammasome to drive caspase-1-dependent IL-1β and IL-18 maturation (Kayagaki et al., 2015; Kayagaki et al., 2011). Consistent with the chemical diversity of NLRP3 stimuli, activation of the NLRP3 inflammasome is thought to converge on sensing of a secondary messenger or cellular state that is universally induced by NLRP3-activating agents (Munoz-Planillo et al., 2013).

Aberrant NLRP3 inflammasome activity is thought to contribute to the pathogenesis of many chronic diseases, including inflammatory diseases such as gout and pseudogout, metabolic diseases like atherosclerosis and NAFLD/NASH, and neurodegenerative diseases like Alzheimer’s disease, Parkinson’s disease and multiple sclerosis (Mangan et al., 2018; Voet et al., 2019). Moreover, gain-of-function mutations in and around the central NACHT domain of NLRP3 cause three autosomal dominantly inherited periodic fever syndromes that together are known as cryopyrin-associated periodic syndrome (CAPS). Symptoms span a clinical spectrum with Familial Cold Autoinflammatory Syndrome (FCAS) being the mildest; Muckle-well syndrome (MWS) being of moderate severity; and Neonatal Onset Multisystem Inflammatory Disease (NOMID)/Chronic Infantile Neurological, Cutaneous and Articular Syndrome (CINCA) being the most severe form of CAPS, featuring systemic inflammation, neurological and sensory impairment and deforming arthropathy (Van Gorp et al., 2019).

Selective and potent inhibitors of the NLRP3 inflammasome may have broad therapeutic potential in CAPS and other diseases (Mangan et al., 2018; Voet et al., 2019). Early studies with the sulfonylurea drug glyburide provided proof-of-concept that PAMP-, DAMP-, and crystal-induced activation of the NLRP3 inflammasome pathway may be selectively targeted without interfering with other inflammasome pathways (Lamkanfi et al., 2009). The related diarylsulfonylurea compound MCC950/CRID3 was originally reported as an inhibitor of IL-1ß secretion (Laliberte et al., 2003), and subsequently shown to potently and selectively inhibit the NLRP3 inflammasome pathway in murine and human macrophages and monocytes with IC_50_ values in the low nM range (Coll et al., 2015; Primiano et al., 2016). There is significant interest in the discovery and development of diarylsulfonylurea CRIDs such as MCC950/CRID3 for the treatment of CAPS and other diseases based on its ability to curb inflammatory pathology in mouse models of CAPS, the experimental autoimmune encephalomyelitis mouse model of multiple sclerosis, NAFLD/NASH, and many other inflammatory disease models (Mangan et al., 2018; Voet et al., 2019). However, the direct molecular target of MCC950/CRID3 in the NLRP3 inflammasome pathway has remains elusive, hampering the rational optimization and development of MCC950/CRID3-based therapies.

Utilizing two different chemoproteomic strategies, we here demonstrate that the NACHT domain of NLRP3 is the molecular target of diarylsulfonylurea CRIDs. Interestingly, we find photoaffinity labelling (PAL) of the NACHT domain of NLRP3 requires an intact (d)ATP-binding pocket and is substantially reduced for most CAPS-associated NLRP3 mutants. In accordance, NLRP3-driven inflammatory pathology in mouse models of CAPS was not efficiently curbed by MCC950/CRID3. Consistently, MCC950/CRID3 abolished circulating levels of interleukin (IL)-1β and IL-18 in LPS-challenged wildtype mice but not in CAPS mice and ex vivo-stimulated mutant macrophages. These results identify the central NACHT domain of wildtype NLRP3 as the molecular target of MCC950/CRID3 and show that CAPS-related NACHT mutations prevent efficient MCC950/CRID3 inhibition. Collectively, this work suggests that MCC950/CRID3-based therapies may effectively treat inflammation driven by wildtype NLRP3, but not CAPS-associated mutants.

## Results

### MCC950/CRID3 selectively binds to human and murine NLRP3

As a first approach to identify the molecular target of MCC950/CRID3, we made use of photo-affinity labeling (PAL) in combination with click chemistry (Smith and Collins, 2015). Guided by limited structure-activity studies, a cell-permeable photo-affinity probe was synthesized (**Fig. 1A**, compound PAL-CRID3) that contains a photo-reactive benzophenone group to enable direct covalent labeling of MCC950/CRID3 targets upon exposure to UV light. The alkyne functionality in PAL-CRID3 allowed in situ click reaction with a 5-carboxytetramethylrhodamine (TAMRA) fluorescent reporter to support in-gel fluorescence detection of the covalent MCC950/CRID3-protein adduct by SDS-PAGE. A dose-response analysis confirmed that PAL- CRID3 retained the ability to potently inhibit NLRP3 inflammasome activation with sub-micromolar IC_50_ values. PAL-CRID3 inhibited nigericin-induced IL-1β secretion from Pam3CSK4-primed primary bone marrow-derived macrophages (BMDMs) (**Fig. 1b**; IC_50_ = 731 nM) and ER-Hoxb8-immortalized macrophages (iMac) (**Fig. 1c**; IC_50_ = 453 nM). As a reference, MCC950/CRID3 inhibited nigericin-induced IL-1β secretion with approximately 8- and 11-fold lower IC_50_ values of 90 and 40 nM, respectively (**Fig. 1b, c**). We hypothesize somewhat lesser activity of PAL-CRID3 compared to MCC950/CRID3 is a combination of physical property and binding differences that decrease target occupancy.

**Figure 1.**
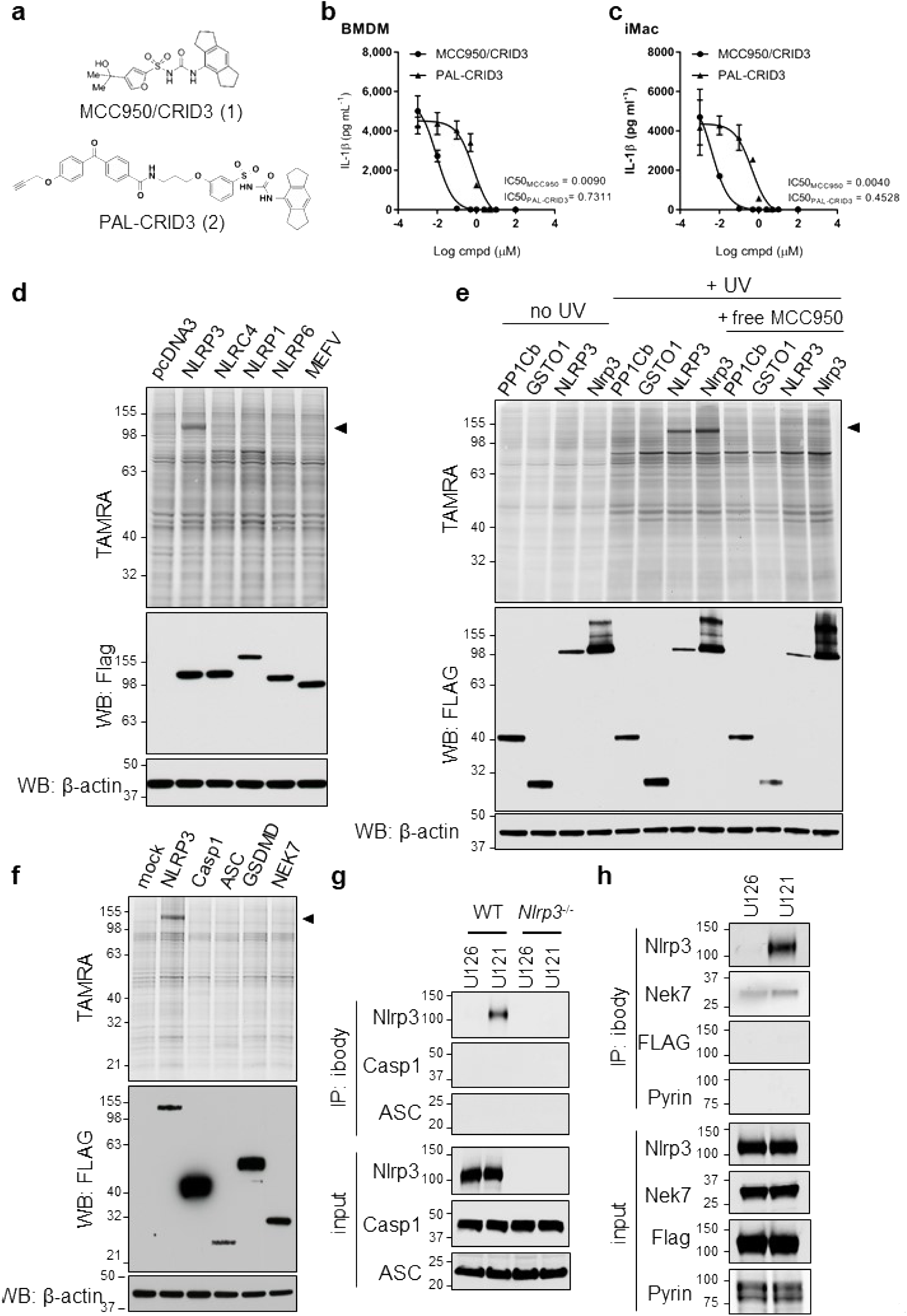
MCC950/CRID3 directly targets NLRP3. (A) Chemical structure of MCC950/CRID3 and the photoaffinity probe PAL-CRID3. (B-C) LPS-primed BMDMs (B) and iMac (C) were stimulated with nigericin in presence of the indicated concentrations of MCC950/CRID3 or PAL-CRID3. Supernatants was analyzed for IL-1β secretion. (D) HEK293 T cells, transfected with the indicated plasmids, were incubated with PAL-CRID3 and subsequently irradiated. After cell lysis and click chemistry conjugation, lysates were assayed by TAMRA imaging and by immunoblot analysis with the indicated antibodies. (E) HEK293 T cells, transfected with the indicated plasmids, were incubated with PAL-CRID3 in absence or presence of free MCC950/CRID3 and subsequently irradiated (+ UV) or left untreated (no UV). After cells are lysed, click chemistry is performed and lysates were assayed by TAMRA imaging and by immunoblot analysis with the indicated antibodies. (F) HEK293 T cells, transfected with the indicated plasmids, were incubated with PAL-CRID3 and subsequently irradiated. After cells are lysed, click chemistry is performed and lysates are assayed by TAMRA imaging and by immunoblot analysis with the indicated antibodies. (G) Lysates of LPS-primed wild type or Nlrp3-deficient BMDMs were incubated with iBody U126 (Ctrl) or iBody U121 (MCC950/CRID3) and analyzed by pull down assay with streptavidin agarose. Precipitates and lysates were assayed by immunoblot analysis with the indicated antibodies. (H) Lysates of LPS- primed Nlrc4^Flag/Flag^ BMDMs were incubated with iBody U126 (Ctrl) or iBody U121 (MCC950/CRID3) and lysates were analyzed by pull down assay with streptavidin agarose. Precipitates and lysates were assayed by immunoblot analysis with the indicated antibodies. Graphs show mean ± s.d. of triplicate wells and represent three independent experiments.

To screen candidate targets of MCC950/CRID3, HEK293 T cells overexpressing FLAG-epitope tagged fusions of the human inflammasome sensor proteins NLRP3, NLRP1, NLRC4, NLRP6 and MEFV were incubated with PAL-CRID3 and exposed to UV light to allow covalent binding of PAL-CRID3 to potential targets. Following cell lysis, the probe was conjugated to the TAMRA reporter by click chemistry, and protein lysates were separated by SDS-PAGE. Notably, in-gel fluorescence imaging showed significant TAMRA labelling of ectopically expressed NLRP3, but not other inflammasome sensors in the panel (**Fig. 1d**). To further extend these findings, we next confirmed binding to murine Nlrp3 (**Fig. 1e**). As controls, protein phosphatase PP1Cb and glutathione S-transferase GSTO1, which has been proposed as the target of CRID compounds (Laliberte et al., 2003), were not labelled by PAL-CRID3 (**Fig. 1e**). Moreover, binding of PAL-CRID3 to human NLRP3 and murine Nlrp3 only was observed following UV crosslinking and labelling was rescued by competition with excess MCC950/CRID3 (**Fig. 1e**), thus ruling out non-specific cross-linking and validating specificity of these findings. To further characterize the interaction of PAL-CRID3 with components of the NLRP3 inflammasome, we assessed binding to human NEK7, ASC, caspase-1 and GSDMD. PAL-CRID3 failed to label the NLRP3 inflammasome components in the panel apart from NLRP3 (**Fig. 1f**). Collectively, these results suggest that CRID compounds, including MCC950/CRID3, inhibit NLRP3 inflammasome signalling by directly binding to NLRP3.

To confirm and extend these findings to endogenous NLRP3, we made use of recently described iBody technology to immobilize MCC950/CRID3 on polymers that enabled immunoprecipitation of MCC950/CRID3 targets (Simon et al., 2018). In brief, MCC950/CRID3 was stochastically modified with a photo-activatable phenyldiazirine linker, and the resulting isomeric mixture was conjugated to a hydrophilic *N*-(2-hydroxypropyl)methacrylamide (HPMA) polymer backbone that is decorated with a biotin affinity tag (for details please refer to the Materials and Methods section and (Simon et al., 2018)). The MCC950/CRID3 iBody conjugate is further referred to as iBody U121. The corresponding iBody conjugate lacking MCC950/CRID3 served as a negative control and is referred to as iBody U126.

BMDMs of wildtype (C57BL/6J) and *Nlrp3^−/−^* mice were primed with LPS to transcriptionally upregulate NLRP3 inflammasome components (Bauernfeind et al., 2009), and cell lysates were subsequently incubated with iBody U121 (MCC950/CRID3) or iBody U126 (control) followed by immunoprecipitation with streptavidin-coupled beads. Contrary to control iBody U126, iBody U121 immunoprecipitated endogenous Nlrp3 from wildtype BMDMs (**Fig. 1g**). iBody U121 failed to pulldown inflammasome components ASC and caspase-1 (**Fig. 1g**). As expected, the immunoreactive band was absent from immunoprecipitates and lysates of LPS-primed *Nlrp3*-deficient macrophages (**Fig. 1g**). Given the lack of suitable antibodies to detect endogenous NLRC4, we made use of reported *Nlrc4*^3xFlag/3xFlag^ knock-in mice (Matusiak et al., 2015; Qu et al., 2012) to further validate selective targeting of NLRP3. Consistent with our earlier results, only iBody U121 precipitated endogenous Nlrp3 from LPS-primed *Nlrc4*^3xFlag/3xFlag^ BMDMs (**Fig. 1h**). Comparable background binding of Nek7 was observed with iBody U121 and control iBody U126, whereas neither Nlrc4 (detected using FLAG antibodies), Nek7 nor Pyrin were retrieved in iBody U121 immunoprecipates of LPS-primed *Nlrc4*^Flag/Flag^ BMDMs (**Fig. 1h**). Collectively, these results show that MCC950/CRID3 selectively binds to NLRP3 in LPS-primed macrophages.

### PAL-CRID3 targets the central NACHT domain of wildtype NLRP3, but not CAPS mutants

NLRP3 consists of an amino-terminal Pyrin domain (PYD), a central nucleotide binding and oligomerization (NACHT) domain and carboxy-terminal leucine-rich repeats (LRRs) that are thought to lock NLRP3 in an inactive conformation. To map the NLRP3 region(s) to which MCC950/CRID3 binds, we generated deletion mutants that cover the following three regions of human NLRP3: (*i*) the N-terminal PYD; (*ii*) the central NACHT domain (comprising the NOD domain and helical domain 2 (HD2)); and (*iii*) the carboxy-terminal LRR region (**Fig. 2a**). Consistent with our previous results (**Fig. 1**), TAMRA analysis showed significant binding of PAL-CRID3 to ectopically expressed full-length NLRP3 in HEK293 T cells (**Fig. 2b**). In addition, we observed binding of PAL-CRID3 to the isolated NACHT (NOD + HD2) region of NLRP3, but not to the PYD and LRR domains (**Fig. 2b**). Labelling of both full-length NLRP3 and the isolated NACHT region by PAL-CRID3 was rescued by treatment with an excess amount of free MCC950/CRID3 (**Fig. 2b**), confirming specificity and suggesting PAL-CRID3 and MCC950/CRID3 compete for the same binding site. Binding of ATP/dATP to the Walker A pocket in the NACHT domain is essential for NLRP3 inflammasome assembly and function (Duncan et al., 2007). Interestingly, mutation of the Walker A motif (GKT^229^/AAA) in full-length Nlrp3 abolished PAL- CRID3 binding (**Fig. 2c**), suggesting that an intact ATP binding pocket is required for strongest binding of diarylsulfonylurea CRID compounds.

**Figure 2.**
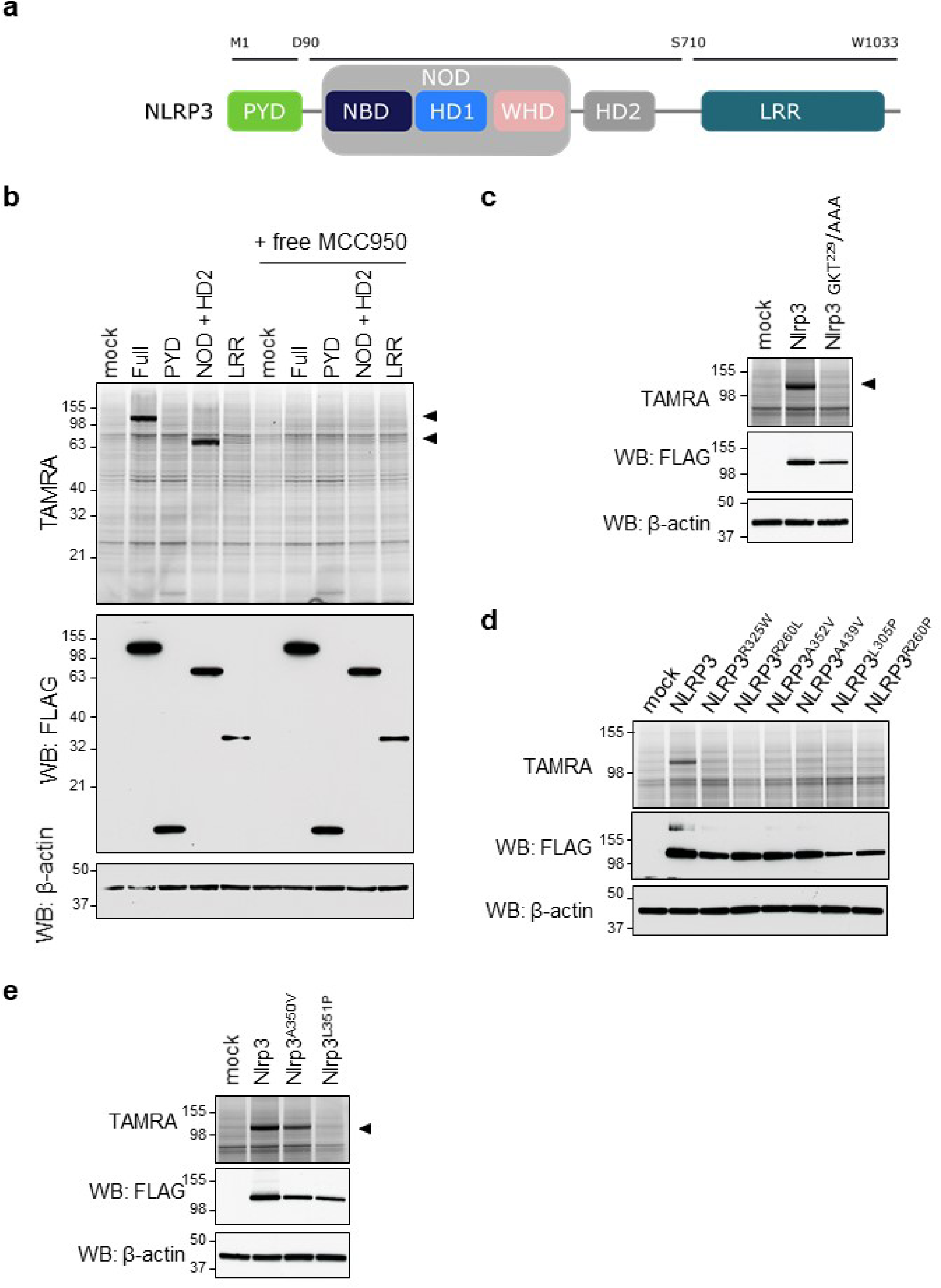
PAL-CRID3 binding requires an intact (d)ATP-binding pocket and fails to bind CAPS- associated NLRP3 mutants. (A) Schematic diagram of NLRP3 depicting the different domains used in this study. PYD, Pyrin domain; NOD, nucleotide-binding oligomerization domain; NBS, nucleotide-binding domain; HD1 and HD2, helical domains 1 and 2; WHD, winged helix domain; LRR, leucine-rich repeat. (B) HEK293 T cells, transfected with the indicated plasmids, were incubated with PAL-CRID3 in absence or presence of free MCC950/CRID3 and subsequently irradiated. After cells are lysed, click chemistry is performed and lysates are assayed by TAMRA imaging and by immunoblot analysis with the indicated antibodies. (C-E) HEK293 T cells, transfected with the indicated plasmids, were incubated with PAL-CRID3 and subsequently irradiated. After cells are lysed, click chemistry is performed and lysates are assayed by TAMRA imaging and by immunoblot analysis with the indicated antibodies.

The central NACHT domain of NLRP3 also contains most reported gain-of-function mutations that cause CAPS disease (https://infevers.umai-montpellier.fr/web/index.php). This prompted us to explore PAL-CRID3 binding to CAPS-associated NLRP3 mutants. Unexpectedly, introducing a random selection of 6 different CAPS mutations in the NACHT domain of human NLRP3 that are associated with respectively MWS (NLRP3^R325W^, NLRP3^R260L^, NLRP3^A352V^), FCAS (NLRP3^A439V^, NLRP3^L305P^) or NOMID (NLRP3^R260P^) all blunted binding of PAL-CRID3 to full-length human NLRP3 (**Fig. 2d**). The classical MWS A352 V and FCAS L353P mutations in human NLRP3 correspond to the A350 V and L351P mutations that have been knocked into the murine *Nlrp3* gene to model CAPS disease in mice (Brydges et al., 2009). Notably, the Nlrp3^A350V^ mutant did not significantly affect labelling by PAL-CRID3, whereas the Nlrp3^L351P^ mutation abolished labelling by PAL-CRID3 (**Fig. 2e**). Together, these results suggest that photoaffinity labelling of the NACHT domain requires an intact (d)ATP-binding pocket and is substantially reduced for most CAPS-associated NLRP3 mutants.

### MCC950/CRID3 inhibits the inflammasome in wildtype but not *Nlrp3*^L351P^ macrophages

Considering that our observation that CAPS-associated Nlrp3 mutants escape PAL-CRID3 binding may have potential implications for treating CAPS with MCC950/CRID3, we sought to functionally validate these results with MCC950/CRID3 in the reported *Nlrp3*^L351P^ CAPS model (Brydges et al., 2009). To this end, mice that were homozygous for the *Nlrp3*^L351P^ allele were bred to transgenic mice that hemizygously expressed the tamoxifen-inducible Cre-ERT2 fusion gene (CreT) (Ventura et al., 2007). Following tamoxifen treatment and excision of the floxed neomycin resistance cassette, BMDMs of the resulting *Nlrp3*^L351P/+^CreT^+^ mice expressed Nlrp3 from both the wildtype and *Nlrp3*^L351P^ alleles. Macrophages from Cre-ERT2-negative littermates (*Nlrp3*^L351P/+^CreT^−^), which only express Nlrp3 from the wildtype allele, were used as controls in these experiments.

Culture media of LPS-stimulated *Nlrp3*^L351P/+^CreT^+^ macrophages contained significant levels of IL-1β (**Fig. 3a**) and IL-18 (**Fig. 3b**) that were associated with marked maturation of caspase-1 and IL-1β in cell lysates (**Fig. 3c**). As reported (Brydges et al., 2009), these LPS-induced inflammasome responses were driven by the CAPS-associated *Nlrp3*^L351P^ allele because *Nlrp3*^L351P/+^CreT^−^ BMDMs failed to secrete detectable levels of IL-1β (**Fig. 3a**) and IL-18 (**Fig. 3b**), and did not contain mature caspase-1 and IL-1β in their cell lysates (**Fig. 3c**). Consistent with our previous results that CAPS-associated Nlrp3^L351P^ escaped PAL-CRID3 binding, we found that MCC950/CRID3 failed to inhibit the above inflammasome responses driven by the *Nlrp3*^L351P^ allele. We confirmed that MCC950/CRID3 was active against wildtype NLRP3 because it abolished levels of LPS-ATP-induced secretion of IL-1β (**Fig. 3d**) and IL-18 (**Fig. 3e**) in culture media of *Nlrp3*^L351P/+^CreT^−^ BMDMs (that are solely driven by wildtype Nlrp3 in this genotype given the absence of a Cre-ERT2 transgene), as well as the concomitant maturation of caspase-1 and IL-1β in the corresponding cell lysates (**Fig. 3f**). Similarly, MCC950/CRID3 abolished LPS+nigericin-induced secretion of IL-1β (**Fig. 3g**) and IL-18 (**Fig. 3h**), and maturation of caspase-1 and IL-1β by *Nlrp3*^L351P/+^CreT^−^ macrophages (**Fig. 3i**). In marked contrast, LPS+ATP- and LPS+nigericin-induced inflammasome responses were insensitive to MCC950/CRID3 blockade in *Nlrp3*^L351P/+^CreT^+^ BMDMs that express both the *Nlrp3*^L351P^ alleleand wildtype Nlrp3 (**Fig. 3d-i**). Together, these results establish that the CAPS-associated *Nlrp3*^L351P^ allele is insensitive to MCC950/CRID3 inhibition.

**Figure 3.**
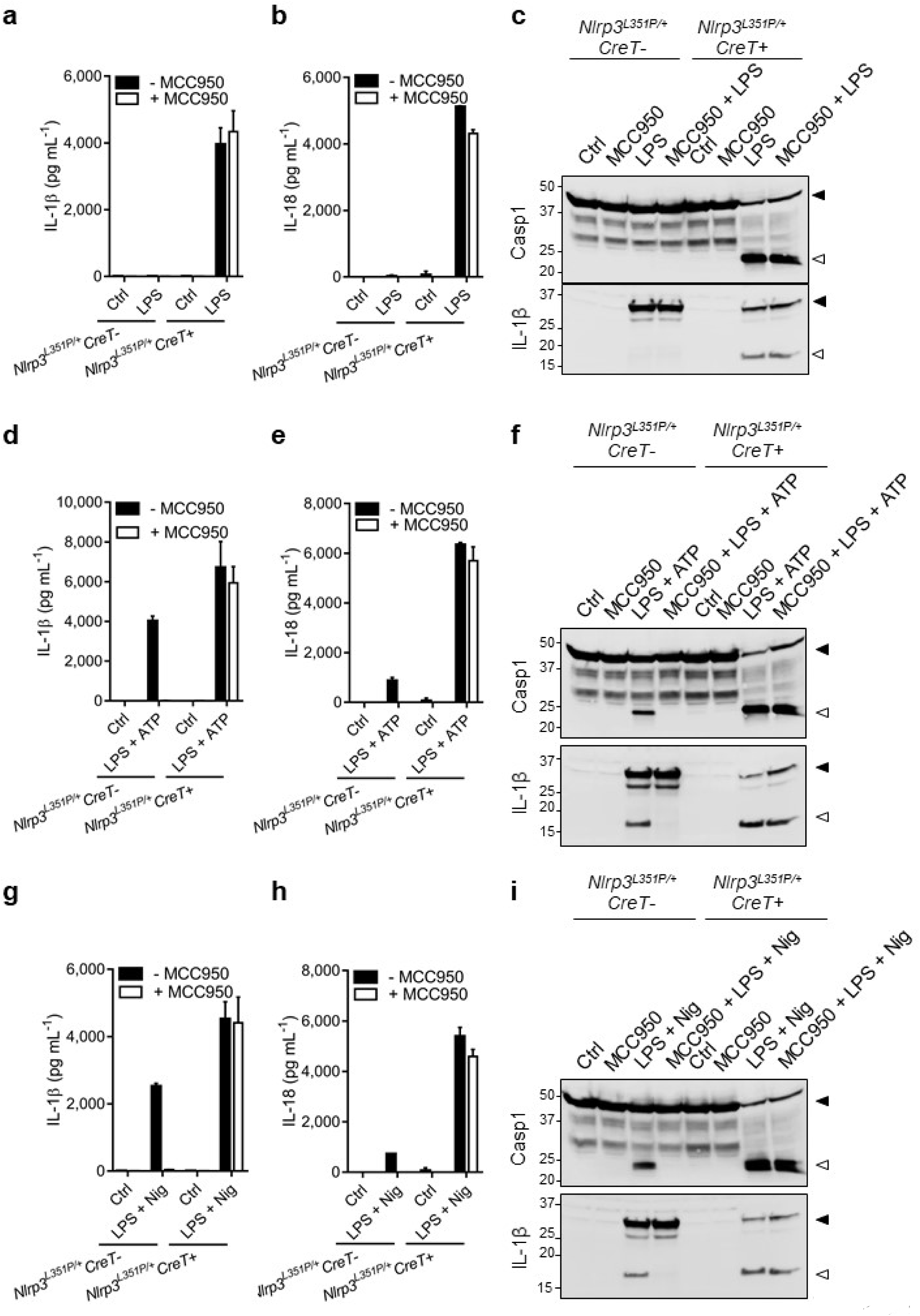
MCC950/CRID3 inhibits the inflammasome in wildtype but not Nlrp3^L351P^ macrophages. (A-C) BMDMs from tamoxifen-treated *Nlrp3^L351P/+^ CreT-* and *Nlrp3^L351P/+^ CreT+* mice were left untreated (Ctrl) or treated with LPS. Supernatants was analyzed for IL-1β (A) and IL-18 (B) secretion and lysates were immunoblotted for caspase-1 and IL-1β (C). (D-F) BMDMs from tamoxifen-treated *Nlrp3^L351P/+^ CreT-* and *Nlrp3^L351P/+^ CreT+* mice were left untreated (Ctrl) or primed with LPS and stimulated with ATP. Supernatants was analyzed for IL-1β (D) and IL-18 (E) secretion and lysates were immunoblotted for caspase-1 and IL-1β (F). (G-I) BMDMs from tamoxifen-treated *Nlrp3^L351P/+^ CreT-* and *Nlrp3^L351P/+^ CreT+* mice were left untreated (Ctrl) or primed with LPS and stimulated with nigericin (Nig). Supernatants was analyzed for IL-1β (G) and IL-18 (H) secretion and lysates were immunoblotted for caspase-1 and IL-1β (I). Graphs show mean ± s.d. of triplicate wells and represent three independent experiments.

### MCC950/CRID3 inhibition of inflammasome responses in *Nlrp3*^A350V^ macrophages

Unlike for Nlrp3^L351P^, labelling of Nlrp3^A350V^ by G03086997 was not significantly impacted by the mutation (**Fig. 2e**). To address whether this was mirrored by potent inhibition of *Nlrp3*^A350V^-driven inflammasome responses by MCC950/CRID3, mice that were homozygous for the previously reported *Nlrp3*^A350V^ allele were bred to the Cre-ERT2 (CreT) transgenic mice described above. Following tamoxifen treatment and excision of the floxed neomycin resistance cassette, BMDMs of the resulting Nlrp3^A350V/+^CreT^+^ mice expressed Nlrp3 from both the wildtype and *Nlrp3*^A350V^ alleles. BMDMs from CreT-negative littermates (*Nlrp3*^A350V/+^CreT^−^), which only express Nlrp3 from the wildtype allele, were used as controls in these experiments.

Like *Nlrp3*^L351P/+^CreT^+^ BMDMs (**Fig. 3a-c**), *Nlrp3*^A350V/+^CreT^+^ macrophages that expressed Nlrp3^A350V^ secreted high levels of secreted IL-1β (**Fig. 4a**) and IL-18 (**Fig. 4b**) in response to LPS stimulation alone. This was accompanied by proteolytic maturation of caspase-1 and proIL-1β as demonstrated by immunoblot analysis (**Fig. 4c**). As expected, these responses were absent from LPS-stimulated *Nlrp3*^A350V/+^CreT^−^ BMDMs that expressed wildtype Nlrp3 only (**Fig. 4a-c**). MCC950/CRID3 potently inhibited LPS-induced IL-1β (**Fig. 4a**) and IL-18 (**Fig. 4b**) secretion, and Nlrp3^A350V^-driven cleavage of caspase-1 and IL-1β (**Fig. 4c**) in marked contrast to *Nlrp3*^L351P/+^CreT^+^ macrophages that proved insensitive to MCC950/CRID3 blockade (**Fig. 3a-c**).

**Figure 4.**
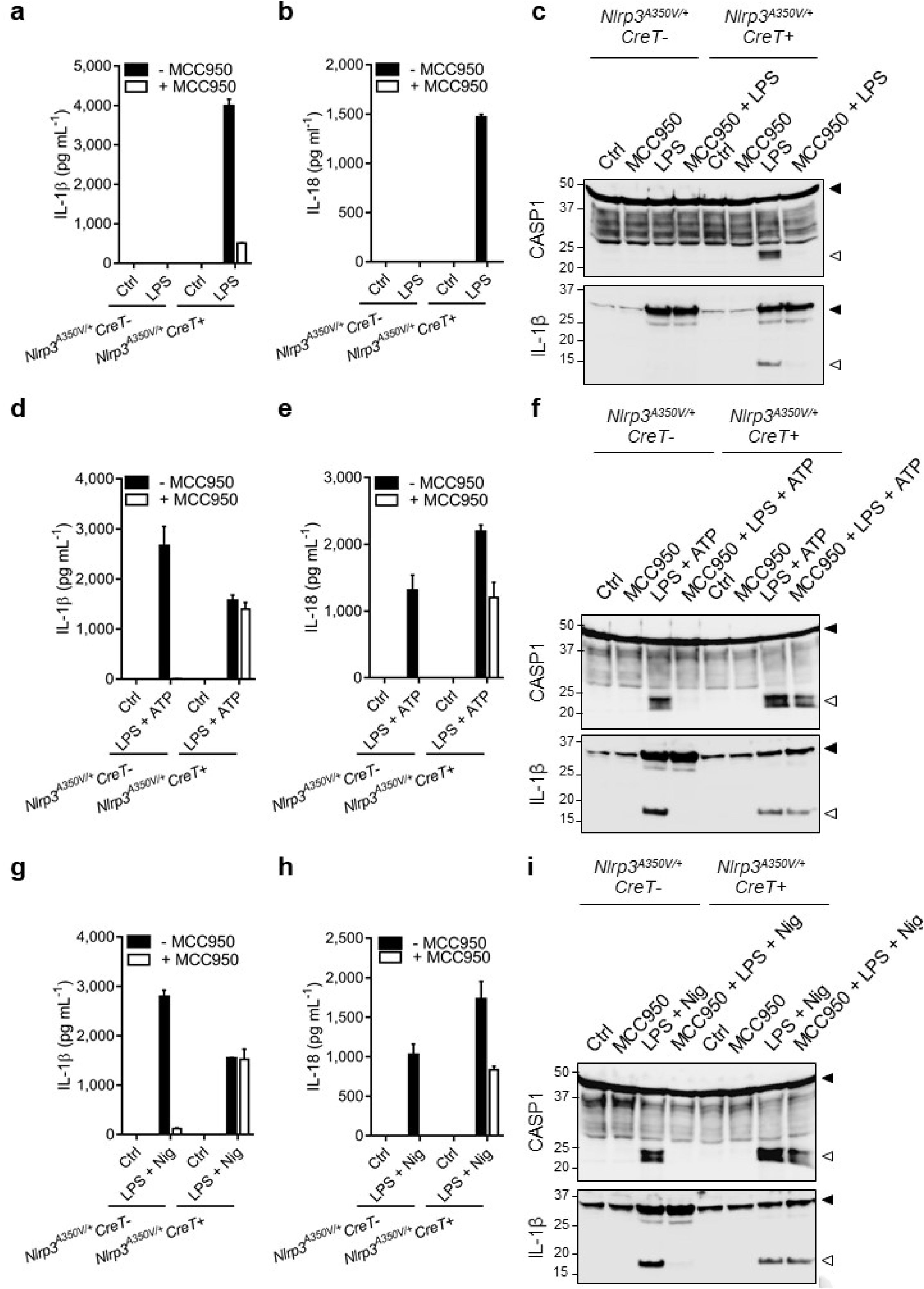
MCC950/CRID3 inhibition of inflammasome responses in Nlrp3^A350V^ macrophages. (A-C) BMDMs from tamoxifen-treated *Nlrp3^A350V/+^ CreT-* and *Nlrp3^A350V/+^ CreT+* mice were left untreated (Ctrl) or treated with LPS. Supernatants was analyzed for IL-1β (A) and IL-18 (B) secretion and lysates were immunoblotted for caspase-1 and IL-1β (C). (D-F) BMDMs from tamoxifen-treated *Nlrp3^A350V/+^ CreT-* and *Nlrp3^A350V/+^ CreT+* mice were left untreated (Ctrl) or primed with LPS and stimulated with ATP. Supernatants was analyzed for IL-1β (D) and IL-18 (E) secretion and lysates were immunoblotted for caspase-1 and IL-1β (F). (G-I) BMDMs from tamoxifen-treated *Nlrp3^A350V/+^ CreT-* and *Nlrp3^A350V/+^ CreT+* mice were left untreated (Ctrl) or primed with LPS and stimulated with nigericin (Nig). Supernatants was analyzed for IL-1β (G) and IL-18 (H) secretion and lysates were immunoblotted for caspase-1 and IL-1β (I). Graphs show mean ± s.d. of triplicate wells and represent three independent experiments.

LPS+ATP and LPS+nigericin potently triggered IL-1β and IL-18 secretion, and maturation of caspase-1 and proIL-1β from both *Nlrp3*^A350V/+^CreT^+^ and *Nlrp3*^A350V/+^CreT^−^ control BMDMs (**Fig. 4d-i**). However, whereas MCC950/CRID3 abolished LPS+ATP- and LPS+nigericin-induced IL-1β secretion from *Nlrp3*^A350V/+^CreT^−^ control BMDMs, it failed to alter secretion of IL-1β from parallelly-treated *Nlrp3*^A350V/+^CreT^+^ BMDMs (**Fig. 4d, g**). MCC950/CRID3 also abrogated LPS+ATP- and LPS+nigericin-induced IL-18 secretion from control *Nlrp3*^A350V/+^CreT^−^ BMDMs, but only reduced IL-18 secretion from parallelly stimulated *Nlrp3*^A350V/+^CreT^+^ macrophages by about 50% (**Fig. 4e, h**). Aligned with these results, MCC950/CRID3 inhibited LPS+ATP- and LPS+nigericin-induced cleavage of caspase-1 and proIL-1β in control *Nlrp3*^A350V/+^CreT^−^ BMDMs but not in *Nlrp3*^A350V/+^CreT^+^ macrophages (**Fig. 4f, i**). Together, these results show that although binding of PAL-CRID3 to Nlrp3^A350V^ was not significantly compromised, the mutation subtly alters the ability of MCC950/CRID3 to inhibit inflammasome activation in primary macrophages, with potent inhibition seen only in response to LPS but not following ‘signal 2′ triggers such as ATP and nigericin.

### Inflammasome inhibition in homozygous mutant macrophages

The studies described above were performed in heterozygous macrophages that express both wildtype and CAPS-associated Nlrp3 mutants. Considering that the Nlrp3 NACHT region facilitates Nlrp3 oligomerization, we decided to further assess MCC950/CRID3 responses in macrophages that uniquely express the CAPS-associated Nlrp3 mutants in the absence of wildtype Nlrp3. To do so, we transduced wildtype, *Nlrp3*^A350V/A350V^ and *Nlrp3*^L351P/L351P^ BMDMs with Cre recombinase-expressing lentiviruses to excise the neomycin resistance cassette that is placed upstream of the Nlrp3 mutation and allow expression of the CAPS-associated Nlrp3 mutants.

As expected, wildtype macrophages failed to secrete IL-1β and IL-18 in response to LPS alone, and MCC950/CRID3 abolished secretion of IL-1β and IL-18 when wildtype BMDM were stimulated with LPS+ATP or LPS+nigericin (**Fig. 5a, b**). LPS stimulation alone was sufficient to induce extracellular release of IL-1β and IL-18 in homozygous *Nlrp3*^L351P/ L351P^ and *Nlrp3*^A350V/ A350V^ macrophages (**Fig. 5c-f**). MCC950/CRID3 inhibited Nlrp3^L351P^-induced secretion of IL-1β and IL-18 neither in response to LPS alone, nor when combined with ‘signal 2′ agents ATP or nigericin (**Fig. 5c, d**), unequivocally establishing that Nlrp3^L351P^-induced inflammasome activation is insensitive to MCC950/CRID3 blockade. Consistent with our previous results in heterozygous *Nlrp3*^A350V^ mutant macrophages, MCC950/CRID3 partially inhibited LPS-induced IL-1β and IL-18 levels in culture supernatants of homozygous *Nlrp3*^A350V/ A350V^ macrophages, but this was not observed in response to LPS+ATP and LPS+nigericin (**Fig. 5e, f**).

**Figure 5.**
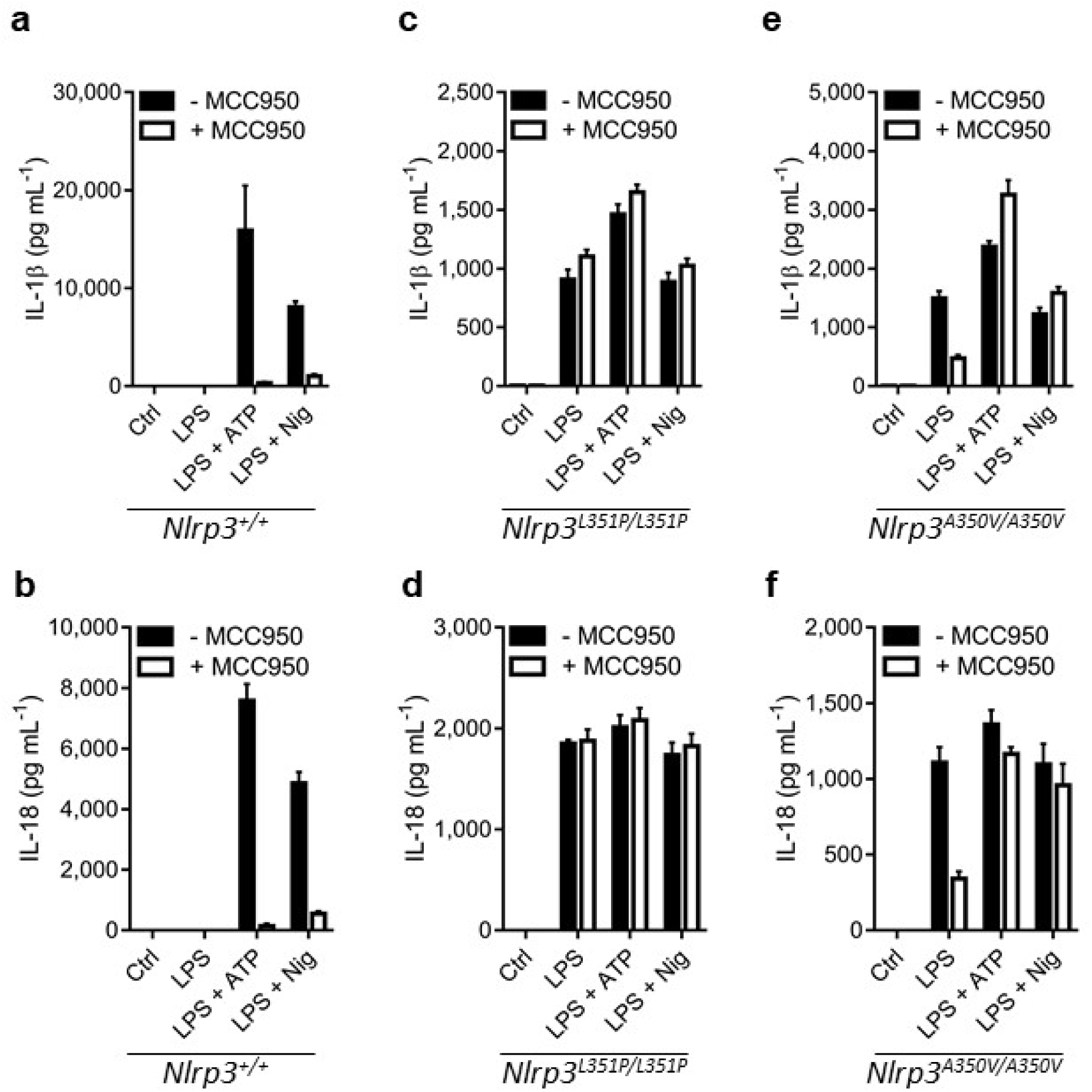
Inflammasome inhibition in homozygous mutant macrophages. (A-F) Wild type (A- B), *Nlrp3^L351P/L351P^* (C-D) or *Nlrp3^A350V/A350V^* (E-F) BMDMs that were transduced with a Cre-recombinase expressing lentiviral vector, were left unstimulated (Ctrl) or stimulated with LPS and then left untreated or treated with ATP or nigericin (Nig) in absence or presence of MCC950/CRID3. Supernatants was analyzed for IL-1β (A,C,E) and IL-18 (B,D,F) secretion. Graphs show mean ± s.d. of triplicate wells and represent three independent experiments.

### *In vivo* MCC950/CRID3 inhibition of inflammasome activation in CAPS disease models

Myeloid-specific expression of the *Nlrp3*^L351P^ and *Nlrp3*^A350V^ alleles in knock-in mice was shown to drive systemic inflammation accompanied by respectively embryonic and perinatal lethality that in both cases required Nlrp3 inflammasome activation (Brydges et al., 2009). To seek further validation of the notion that the *Nlrp3*^L351P^ mutations escapes MCC950/CRID3 inhibition, we next investigated how MCC950/CRID3 treatment impacts on the CAPS phenotype of *Nlrp3*^L351P/+^CreT^+^ mice. As expected, serum levels of IL-1β, IL-18 and IL-6 were significantly increased in *Nlrp3*^L351P/+^CreT^+^ mice three days after tamoxifen dosing relative to the basal levels of tamoxifen-treated *Nlrp3*^L351P/+^CreT^−^ littermate mice (**Supplementary Fig. 1a-c**). Moreover, tamoxifen treatment resulted in *Nlrp3*^L351P/+^CreT^+^ mice presenting with substantial weight loss and mortality, with all mice being lost or requiring termination because of humane endpoints within 5 days after commencing tamoxifen treatment (**Fig. 6a, b**). As a control, tamoxifen administration did not alter body weight or survival of *Nlrp3*^L351P/+^CreT^−^ mice (**Fig. 6a, b**). Daily *i.p.* injection of MCC950/CRID3 did not rescue body weight loss or mortality rates of *Nlrp3*^L351P/+^CreT^+^ mice, suggesting that MCC950/CRID3 failed to inhibit *in vivo* Nlrp3^L351P^-induced inflammasome activation (**Fig. 6a, b**). In agreement, *i.p.* dosing of MCC950/CRID3 failed to reduce tamoxifen-induced levels of IL-1β, IL-18 and IL-6 in serum of *Nlrp3*^L351P/+^ CreT^+^ mice (**Fig. 6c-e**). Thus, the CAPS-associated L351P mutation renders Nlrp3 insensitive to MCC950/CRID3 inhibition.

**Figure 6.**
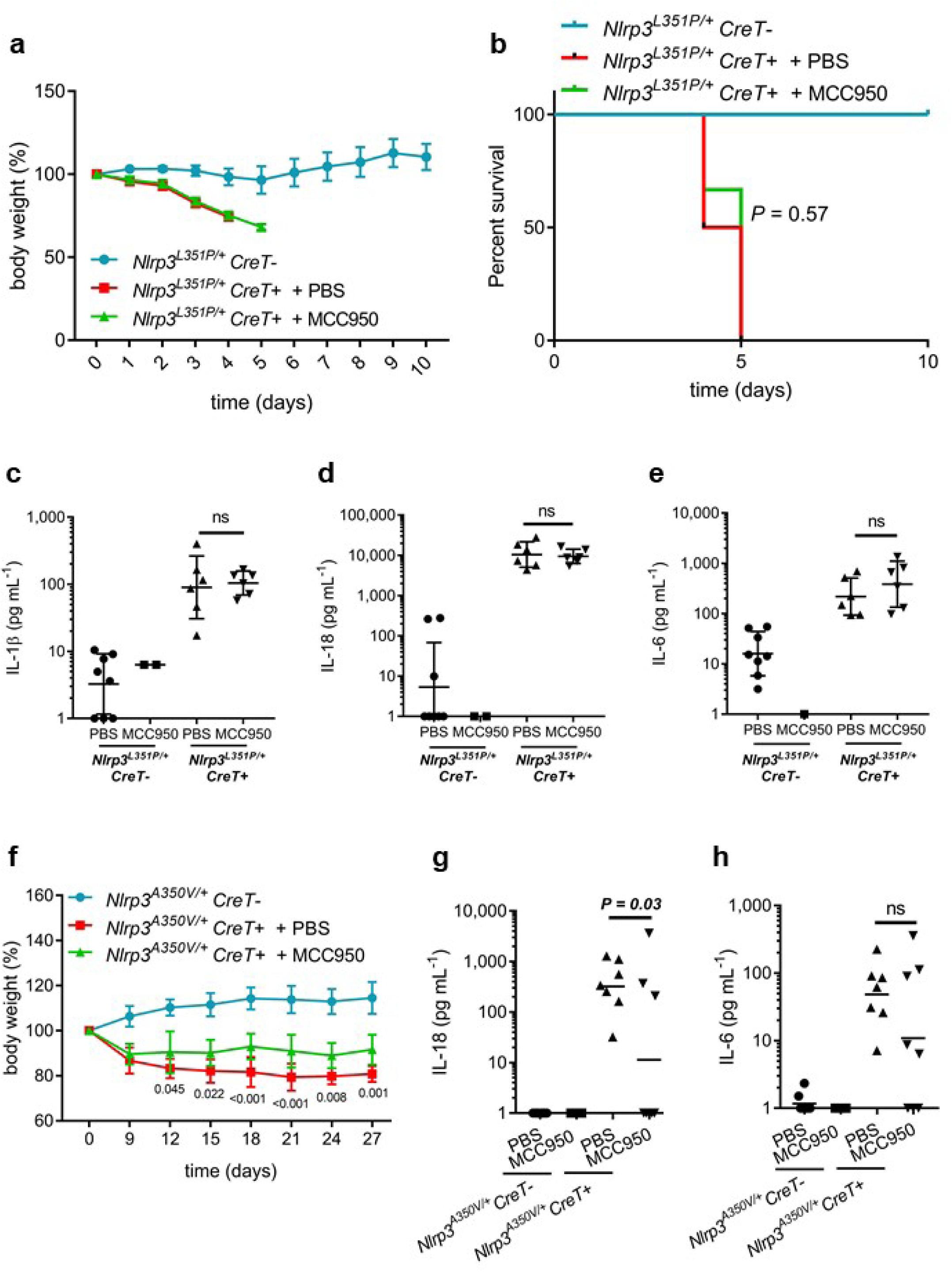
*In vivo* MCC950/CRID3 inhibition of inflammasome activation in CAPS disease models. (A-B) Growth (A) and survival (B) curves of tamoxifen-treated *Nlrp3^L351P/+^ CreT+* that received daily injections of PBS (n = 6) or 50 mg kg^−1^ MCC950/CRID3 (n = 6) and of tamoxifen-treated *CreT-* littermates (n = 6). (C-E) IL-1β (C), IL-18 (D) and IL-6 (E) cytokine levels in serum obtained at day 4 from tamoxifen-treated *Nlrp3^A350V/+^ CreT+* mice that received daily injections of PBS (n = 6) or MCC950/CRID3 (n = 6). Serum of tamoxifen-treated *CreT-* littermates injected with PBS (n = 8) or MCC950/CRID3 (n = 2) was used as control. (F) Growth curves of tamoxifen-treated *Nlrp3^A350V/+^ CreT+* that received daily injections of PBS (n = 6) or 50 mg kg^−1^ MCC950/CRID3 (n = 7) and of tamoxifen-treated *CreT-* littermates (n = 6). (G-H) IL-18 (G) and IL-6 (H) cytokine analysis of serum obtained at day 4 from tamoxifen-treated *Nlrp3^A350V/+^ CreT+* mice that received daily injections of PBS (n = 7) or MCC950/CRID3 (n = 8). Serum of tamoxifen-treated *CreT-* littermates treated with PBS (n = 8) or MCC950/CRID3 (n = 5) was used as control. Graphs show mean ± s.d. and represent two independent experiments.

Our results in *Nlrp3*^A350V^ macrophages showed that this mutation subtly altered the ability of MCC950/CRID3 to inhibit inflammasome activation with potent inhibition seen only in response to LPS but not following ‘signal 2’ triggers such as ATP and nigericin (**Fig. 4**). To determine how this translates to the *in vivo* disease setting, we analyzed circulating cytokine levels and body weight loss of *Nlrp3*^A350V/+^CreT^+^ mice following tamoxifen administration. Although there was a trend towards increased serum levels of IL-1β, the low measured concentrations did not reach statistical significance compared to tamoxifen-treated *Nlrp3*^A350V/+^CreT^−^ littermate mice (**Supplementary Fig. 1d**). However, serum concentrations of IL-18 and IL-6 were signficantly elevated in tamoxifen-treated *Nlrp3*^A350V/+^CreT^+^ mice relative to *Nlrp3*^A350V/+^CreT^−^ littermates (**Supplementary Fig. 1e, f**). Nevertheless, levels of the latter cytokines were on average 10 to 20-fold lower than seen in tamoxifen-treated *Nlrp3*^L351P/+^CreT^+^ mice, a finding that is consistent with the milder pathology and the lack of mortality associated with the *Nlrp3*^A350V/+^CreT^+^ CAPS model. Consistent with published findings (McGeough et al., 2012), *Nlrp3*^A350V/+^CreT^+^ mice developed an inflammatory phenotype characterized by a steady weight loss of up to 20% within 27 days (**Fig. 6e**). Notably, daily *i.p.* dosing of MCC950/CRID3 stabilized *Nlrp3*^A350V^-mediated body weight loss in *Nlrp3*^A350V/+^CreT^+^ mice relative to PBS-treated controls, although differences were small and MCC950/CRID3-treated mice failed to thrive and gain weight like the *Nlrp3*^A350V/+^CreT^−^ control group (**Fig. 6e**). In agreement, MCC950/CRID3 had a mild or no effect on circulating levels of the systemic inflammatory markers IL-18 (**Fig. 6f**) and IL-6 (**Fig. 6g**), respectively. Together, these results establish that MCC950/CRID3 has a weak, but measurable effect on Nlrp3^A350V^- induced inflammasomopathy in adult mice.

### MCC950/CRID3 inhibition of LPS-induced cytokines in CAPS mutant mice

To complement the chronic CAPS disease models described above, we next evaluated the potency of MCC950/CRID3 in inhibiting acute Nlrp3-dependent inflammasome responses by subjecting wildtype and CAPS mutant mice to LPS-induced endotoxemia and probing the effect of MCC950/CRID3 on well-documented Nlrp3-dependent readouts such as LPS-induced elevation of serum levels of IL-1β and IL-18 (He et al., 2013).

As expected, *Nlrp3*^L351P/+^CreT^−^ control mice presented with increased serum levels of IL-1β and IL-18 3 hours after LPS challenge (**Fig. 7a, b**). MCC950/CRID3 substantially curbed circulating IL-1β and IL-18 levels in this control group (**Fig. 7a, b**), consistent with reported findings in LPS- challenged C57BL/6 mice (Coll et al., 2015). Contrastingly, serum levels of IL-1β and IL-18 in LPS-dosed *Nlrp3*^L351P/+^CreT^+^ mice were not significantly impacted by MCC950/CRID3 relative to PBS-treated *Nlrp3*^L351P/+^CreT^+^ littermates (**Fig. 7c, d**).

**Figure 7.**
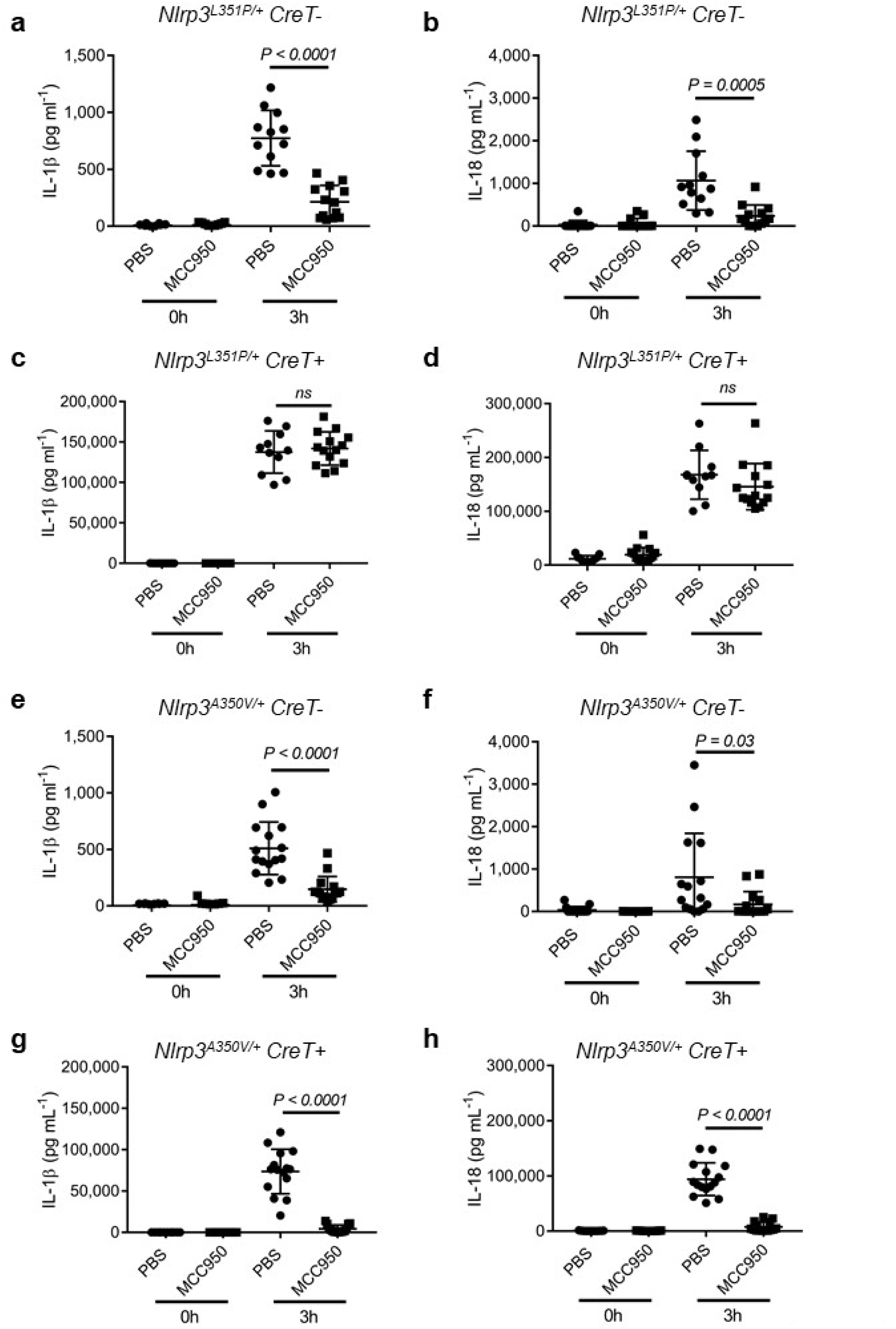
MCC950/CRID3 inhibition of LPS-induced cytokines in CAPS mutant mice. (A-B) At day 3 post-tamoxifen, *Nlrp3^L351P/+^ CreT-* mice were pretreated with PBS (n = 12) or MCC950/CRID3 (n = 13) and subsequently challenged with LPS for 3 hrs. Serum levels of IL-1β (A) and IL-18 (B) were analyzed before and after LPS administration. (C-D) At day 3 post-tamoxifen, *Nlrp3^L351P/+^ CreT+* mice were pretreated with PBS (n = 11) or MCC950/CRID3 (n = 14) and subsequently challenged with LPS for 3 hrs. Serum levels of IL-1β (C) and IL-18 (D) were analyzed before and after LPS administration. (E-F) At day 3 post-tamoxifen, *Nlrp3^A350V/+^ CreT-* mice were pretreated with PBS (n = 15) or MCC950/CRID3 (n = 15) and subsequently challenged with LPS for 3 hrs. Serum levels of IL-1β (E) and IL-18 (F) were analyzed before and after LPS administration. (G-H) At day 3 post-tamoxifen, *Nlrp3^A350V/+^ CreT+* mice were pretreated with PBS (n = 15) or MCC950/CRID3 (n = 15) and subsequently injected with LPS for 3 hrs. Serum levels of IL-1β (G) and IL-18 (H) were analyzed before and after LPS challenge. Graphs show mean ± s.d. and represent two independent experiments.

LPS challenge increased serum concentrations of IL-1β and IL-18 in *Nlrp3*^A350V/+^CreT^−^ control mice, which were inhibited by MCC950/CRID3 (**Fig. 7e, f**) similarly to its effect on LPS-treated *Nlrp3*^L351P/+^CreT^−^ control mice (**Fig. 7a, b**). This is not unexpected since both genotypes express solely wildtype Nlrp3 in the absence of Cre-recombinase-mediated excision of the neomycin resistance cassette that disrupts expression of the respective CAPS-associated Nlrp3 mutants. However, in marked contrast to Nlrp3^L351P^-expressing CreT^+^ mice (**Fig. 7c, d**), MCC950/CRID3 markedly curbed circulating concentrations of IL-1β and IL-18 in Nlrp3^A350V^-expressing CreT^+^ mice (**Fig. 7g, h**). These results confirm that MCC950/CRID3 potently inhibits inflammasome activation by wildtype Nlrp3 and the MWS-associated Nlrp3^A350V^ mutant, but not the FCAS-associated Nlrp3^L351P^ mutant.

## Discussion

Activation of the NLRP3 inflammasome has been observed in many diseases, and its central role in driving pathological inflammation renders it an attractive target for therapeutic intervention (Mangan et al., 2018). Gain-of-function mutations in NLRP3 cause hereditary periodic fever syndromes that are collectively referred to as CAPS, and these patients are currently treated with biologics that target secreted IL-1 (Van Gorp et al., 2019). As reported (Ridker et al., 2017), chronic IL-1-blockade increases risk for fatal infections and sepsis, suggesting that selective targeting of the NLRP3 inflammasome may potentially be a safer and more efficacious therapeutic strategy as it would block production of the central inflammatory mechanisms that contribute to inflammatory pathology while at the same time keeping non-targeted inflammasomes available to produce IL-1β to cope with infections.

Early studies with the sulfonylurea compound glyburide provided proof-of-concept that small molecules may selectively inhibit the NLRP3 inflammasome pathway without interfering with other inflammasomes (Lamkanfi et al., 2009). Subsequently, several additional compounds have been reported to specifically inhibit the NLRP3 inflammasome pathway, but the majority of these agents has weak activity against the NLRP3 inflammasome pathway (μM IC_50_ concentrations) and may target NF-κB signaling and other immune pathways (reviewed in (Mangan et al., 2018)). MCC950/CRID3 is structurally related to glyburide (Coll et al., 2015; Laliberte et al., 2003) and is considered the most potent and selective inhibitor of NLRP3 inflammasome signaling reported to date. There is substantial interest in developing this chemical scaffold for the treatment of CAPS and other diseases. However, these efforts are constrained by the lack on insight in the molecular target and mechanism by which MCC950/CRID3 and related sulfonylurea molecules inhibit activation of the NLRP3 inflammasome pathway.

Making use of photoaffinity labeling and iBody technology as complementary chemical biology approaches, we here identified NLRP3 as the physical target of MCC950/CRID3. We further mapped the binding pocket to the central NACHT domain of NLRP3 and showed that photoaffinity labelling required an intact ATP/dATP binding pocket. This suggests that binding may occur at the nucleotide-binding pocket of NLRP3, although it cannot be excluded that mutations in the Walker A motif may cause long-distance conformational changes that distort a remote MCC950/CRID3 binding pocket elsewhere in the NACHT domain. Notable in this regard is our observation that a randomly selected panel of 6 CAPS-associated gain-of-function mutations in human NLRP3 all failed to be labelled by our PAL probe, further supporting the notion that the sulfonylurea CRID binding pocket is highly sensitive for conformational changes in the protein that are imposed by mutations. Further insight in the conformational requirements and the molecular mechanism by which MCC950/CRID3 inhibits NLRP3 activation awaits a high-resolution structural analysis of the MCC950/CRID3 binding pocket.

Considering the potential implications of these findings for treating CAPS with MCC950/CRID3- based therapies, we evaluated the functional impact of MCC950/CRID3 in two reported CAPS models (Brydges et al., 2009). When knocked into the murine Nlrp3 sequence, the FCAS-associated Nlrp3^L351P^ mutant (corresponding to human L353P) also failed to bind to PAL-CRID3. Consistent with these results, our analysis of both ex vivo-stimulated mutant macrophages and in vivo CAPS and endotoxemia models unequivocally established that MCC950/CRID3 potently inhibits inflammasome activation by wildtype Nlrp3, but not the FCAS-associated Nlrp3^L351P^ mutant. Surprisingly, however, we found that labelling by PAL-CRID3 to the MWS- linked Nlrp3^A350V^ mutant (corresponding to adjacent residue A352 V in human NLRP3) was not significantly impacted by the mutation. MCC950/CRID3 partially inhibited LPS-induced IL-1β and IL-18 secretion from macrophages that homozygously or heterozygously expressed the Nlrp3^A350V^ mutant, but inhibition of the Nlrp3^A350V^ inflammasome was completely lost in response to ‘signal 2’ agents ATP and nigericin. Consequently, MCC950/CRID3 significantly lowered IL-1β and IL-18 levels in serum of LPS-challenged *Nlrp3*^A350V^ knock-in mice, whereas it provided only limited protection against chronic CAPS mutation-driven body weight loss. The CreERT2 recombinase expression system used with our CAPS disease models allows for controlled ubiquitious tissue expression of the mutant Nlrp3 knock-in allele upon tamoxifen treatment in adult animals. Expression of the *Nlrp3*^A350V^ allele does not induce lethality in adult mice under these conditions. However, another study (Coll et al., 2015) that relied on Lysosome M-Cre-driven expression of the *Nlrp3*^A350V^ allele in cells of the myeloid lineage observed that MCC950/CRID3 rescued neonatal lethality, consistent with our observation that the Nlrp3^A350V^ mutant retained sensitivity to MCC950/CRID3 inhibition. This report also suggested MCC950/CRID3 inhibits LPS-induced IL-1β processing in peripheral blood mononuclear cells (PBMCs) from MWS patients. Another study showed that unlike samples from healthy controls, LPS- and LPS+ATP-induced IL-1β secretion from whole blood samples of genetically defined CAPS patients resisted MCC950/CRID3 inhibition (Grinstein et al., 2018), which is consistent with our results suggesting that MCC950/CRID3-based therapies may effectively treat inflammation driven by wildtype NLRP3, but may be less effective in CAPS patients. To conclude, by identifying the molecular target of MCC950/CRID3 in the NLRP3 inflammasome pathway, and by evaluating its ability to inhibit CAPS mutatant variants, the findings presented here provide a mechanistic framework for advancing therapeutic development of this chemical scaffold and for understanding its therapeutic potential in patients.

## Material and methods

### Synthesis of PAL-CRID3

Detailed experimental procedures for the synthesis of PAL-CRID3 are provided in the Supplementary Data section.

### Synthesis of iBody conjugates

Experimental procedures for the preparation of the iBody conjugates have been described by (Simon et al., 2018) and details for the production of MCC950/CRID3 iBody conjugates are provided in the Supplementary Data section.

### iMac and BMDM culture

Bone marrow cells from C57BL/6 mice were immortalized by ER- Hoxb8 (iMac) as described previously (Wang et al., 2006). Primary bone marrow or iMac progenitor cells were differentiated in DMEM or IMDM supplemented with 10% endotoxin-free heat-inactivated fetal bovine serum, 20-30% L929-conditioned medium, 100 U ml^−1^ penicillin, and 100 mg ml^−1^ streptomycin for 5-6 days at 37°C in a humidified atmosphere containing 5% CO_2_. 6 days later, cells were collected and seeded at a density of 8.5×10^5^ cells per well in 12-well plates in IMDM containing 10% heat-inactivated FBS and 1% non-essential amino acids in the presence of antibiotics. The next day BMDMs were either left untreated or treated with 1 μM MCC950/CRID3 (S7809, Selleckchem) and then stimulated with 0.5 μg ml^−1^ ultrapure LPS from *Salmonella minnesota* (tlrl-smlps, Invivogen) for 3 h followed by 5 mM ATP (10519987001, Roche) or 20 μM nigericin (N-7143, Sigma-Aldrich) for 45 min. For the dose-response analysis of PAL-CRID3 and MCC950/CRID3, adherent BMDMs or differentiated iMac were seeded at 1×10^5^ cells per well in 96-well plates and cultured overnight. The following day, medium was removed and replaced with OPTI-MEM I (Thermo Fisher Scientific) containing 1 μg mL^−1^ Pam3CSK4 (Invivogen). Post-priming (5-6 h later), cells were incubated with DMSO (1:1,000), MCC950/CRID3 (0.001-100 µM), or PAL-CRID3 (0.001-100 µM) for 30 min and then stimulated with 5 μg mL^−1^ nigericin (Invivogen) for 30 min.

### Transfections

HEK293 T cells were maintained in DMEM media supplemented with 10% fetal bovine serum. HEK293 T cells were transfected in 12-well plates with Lipofectamine 2000 (Thermo Fisher Scientific) according to manufacturer’s instructions. All constructs, including NLRP3 cDNAs were synthesized and subcloned into pCDNA3 (+) Zeo (Thermo Fisher Scientific).

### Photo-labeling

Transfected HEK293 T were incubated with DMSO or photo probe PAL-CRID3 (1 μM) for 30 min at 37°C in 500 μL OPTI-MEM. For competition experiments with free MCC950/CRID3, 10 μM MCC950/CRID3 was added 30 min prior. Photo-labeling and click chemistry experiments were performed as described with slight modification (Mackinnon and Taunton, 2009). Briefly, cells were washed once with ice-cold PBS and UV irradiated at 365 nm (100 W) on ice for 10 min. Cells were washed once with ice-cold PBS and frozen at −80°C. Lysis buffer was added (40 mM HEPES, 140 mM NaCl, 0.1% Triton-X-100, Roche EDTA-free complete protease inhibitor) and cells were detached with a cell scraper. Crude lysates were clarified by centrifugation (30 min at 14,000 rpm, 4°C). For click chemistry, lysates (20 μL) were incubated with 5 μL freshly mixed click cocktail comprised of: 1.7 mM TBTA in 1:4 DMSO/*t-*BuOH (1.5 μL); 5 mM TAMRA-N_3_ (0.3 μL); 50 mM TCEP (0.5 μL, freshly prepared); 10% SDS (2.7 μL). Then, 50 mM CuSO_4_ (0.5 μL), was added and reactions were incubated for 1 hr at RT with gentle mixing. Reactions were quenched with 4X LDS buffer and separated by SDS-PAGE. Gels were scanned for TAMRA fluorescence on a Typhoon Trio scanner (GE Life Sciences).

### Mice

*Nlrp3^−/−^* (Mariathasan et al., 2006), *NLRC4^3xFlag^ (Qu et al., 2012)* were described. The CAPS models *Nlrp3^A350VneoR^* and *Nlrp3^L351PneoR^* (Brydges et al., 2009), and *R26-Cre^ERT2^* mice (Ventura et al., 2007) (B6.129-*Gt(ROSA)26Sor^tm1(cre/ERT2)Tyj^*/J, Jax stock number: 008463; here abbreviated as CreT) were originally obtained from The Jackson Laboratories and colonies were further maintained at animal facilities of Ghent University. Mice were housed in individually ventilated cages and kept under pathogen-free conditions at animal facilities of Ghent University and Genentech. All animal experiments were conducted with permission of the Ethical committees on laboratory animal welfare of Ghent University. *Nlrp3^A350VneoR^* and *Nlrp3^L351PneoR^* that are homozygous for the mutated Nlrp3 gene were bred to the tamoxifen-inducible Cre line *R26-Cre^ERT2^* mice to generate *Nlrp3^A350neoR/+^ R26-Cre^ERT2^ Tg+* (herein referred to as *Nlrp3^A350V/+^CreT^+^)* and *Nlrp3^L351PneoR/+^R26-Cre^ERT2^ Tg^+^ mice (*here referred to as *Nlrp3^L351P/+^CreT*^+^), in which expression of mutant Nlrp3 is induced through administration of tamoxifen. 4-5 week old mice received at two consecutive days tamoxifen (T5648, Sigma, dissolved in 1:9 ethanol:corn oil (C-8267, Sigma) at 50 mg ml^−1^) through oral gavage at a dose of 5 mg tamoxifen per mouse per day. On day 3, tamoxifen was administered through diet (Teklad Global 16% Rodent Diet met 400 ppm Tamoxifen per kg, Harlan).

### *in vivo* LPS challenge

6-12 weeks old mice were intraperitoneally injected with PBS or 50 mg kg^−1^ MCC950/CRID3, 30 minutes before being challenged with 40 mg kg^−1^ LPS (*E. coli*, serotype 0111:B4, L-2630, Sigma). Mice were euthanized 3 hours after LPS challenge for blood collection. At the 0 h timepoint before LPS challenge, blood was collected by retro-orbital bleeding.

### iBody immunoprecipitation

12×10^6^ BMDMs were harvested and cells were washed in PBS and lysed by three cycles of freeze/thawing in PBS with 0.09% NP40 supplemented with Complete Protease Inhibitor cocktail (4693159001, Roche Applied Science). Cell lysates were clarified by centrifugation at 14.000 rpm for 20 minutes and supernatants was subsequently incubated with 1 μM of the indicated iBody at RT for 10 minutes. Then prewashed streptavidin conjugated beads were added, followed by overnight incubation at 4°C. The next day, beads were washed 3 times in PBS with 0.09% NP40 buffer and biotinylated proteins were eluted in Laemmli buffer and analyzed by Western blot.

### Lentiviral Cre transduction

To induce lentivirus production, the lentiviral GFP.Cre empty vector (20781, Addgene) together with 2^nd^ generation packaging vector psPAX2 (12260, Addgene) and VSV-G-expressing envelope plasmid pCMV-VSV-G (8454, Addgene) were transfected into HEK293 T cells using jetPRIME transfection reagent (114-15, PolyPlus Transfect). Lentiviral particle-containing medium was collected 48 hours after transfection, filtered using a 0.45 μm filter and incubated with harvested bone marrow. 6 days later, differentiated macrophages were collected, washed and seeded into 12-well cell culture plates prior to stimulation.

### Cytokine analysis

Cytokine levels in culture medium and serum were determined by magnetic bead-based multiplex assay using Luminex technology (Bio-Rad), IL-1β ELISA (R&D Systems) and mouse IL-1β tissue culture kit (Meso Scale Discovery) according to the manufacturers’ instructions.

### Western blotting

Cell lysates were prepared using lysis buffer containing 20 mM Tris HCl pH 7.4, 200 mM NaCl and 1% NP-40. Samples for detection of caspase-1 and IL-1β processing were prepared by combining cell lysates with culture supernatants. Samples were denatured in Laemlli buffer and boiled at 95°C for 10 min. SDS-PAGE-separated proteins were transferred to PVDF membranes and immunoblotted with primary antibodies against caspase-1 (AG-20B- 0042-C100, Adipogen), IL-1β (GTX74034, Genetex), Flag-tag (Flag M2-Peroxidase, Sigma-Aldrich) and actin (AC15, Novus Biologicals). Horseradish peroxidase-conjugated goat anti-mouse (115-035-146, Jackson Immunoresearch Laboratories) or anti–rabbit secondary antibody (111-035-144, Jackson Immunoresearch Laboratories) was used to detect proteins by enhanced chemiluminescence (Thermo Scientific).

### Statistical analysis

GraphPad Prism 5.0 software was used for data analysis. For survival studies, data were compared by log rank (Mantel-Cox) test. Two-way ANOVA test were used to assess body weight differences between groups. Unpaired 2-tailed Student’s t-test was applied to compare cytokine serum levels. Data are shown as mean with standard deviation. *P* < 0.05 was considered to indicate statistical significance.

## Acknowledgements

We thank Karen O’Rourke, Aaron Gupta, Nataliya Popovych, Wayne Fairbrother and Victoria Pham (Genentech) for discussion and technical support. We thank Hall Hoffman (UCSD) and Vishva M. Dixit (Genentech) for mutant mice and Tomáš Knedlík (The Czech Academy of Sciences) for help with iBody experiments. This work was supported by grant GA16-02938S from the Grant Agency of the Czech Republic to J.K., and European Research Council Grant 683144 (PyroPop) and the Baillet Latour Medical Research Grant to M.L.

## Author contributions

LVW, IBS, BLL, DD, SW, ARW, PS, JK, NK and ML designed experiments; LVW, IBS, PS, BLL, DD, AF, FVH, PHVS, PS, VS, LK, PeS, SY and SS performed experiments; LVW, IBS, BLL, DD, CES, STS, SY, JK, NK and ML analysed data; PS, PeS, VS, LK, CES, VCP, STS and SY provided essential reagents; LVW, IBS, NK and ML wrote the manuscript with input from all authors; ML oversaw the project.

## Supplementary Materials & Methods

### Synthesis of PAL-CRID3

All reactions were carried out under a nitrogen atmosphere. All commercial reagents and anhydrous solvents were used without additional purification. Nuclear magnetic resonance (NMR) spectra were acquired on a Bruker BioSpin GmbG operating at 400 and 100 MHz for ^1^H and ^13^C, respectively and are referenced internally according to residual solvent signals. NMR data were processed using MNova software and recorded as follows: ^1^H-NMR - chemical shift (δ, ppm), multiplicity (s, singlet; d, doublet; t, triplet; q, quartet; m, multiplet), coupling constant (Hz), and integration; ^13^C-NMR – chemical shift (δ, ppm). High-resolution mass spectra (HRMS) were recorded on a Thermo Scientific Orbitrap Q Exact mass spectrometer. Thin-layer chromatography was performed on EMD TLC Silica gel 60 F254 plates and visualized with UV light. Reactions were monitored by a Shimadzu LCMS/UV system with LC-30AD solvent pump, 2020 MS, Sil-30AC autosampler, SPD-M30A UV detector, CTO-20A column oven, using a 2-98% acetonitrile/0.1% formic acid (or 0.001% ammonia) gradient over 2.5 minutes. Flash column chromatography purifications were done on a Teledyne Isco Combiflash Rf utilizing Silicycle HP columns using a mobile phase composed of either heptane/isopropyl acetate or dichloromethane/methanol.

#### Preparation of tert-Butyl (3-(3-sulfamoylphenoxy)propyl)carbamate

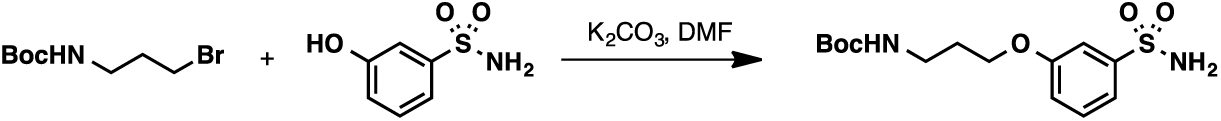

Potassium carbonate (1.00 g, 7.5 mmol) was added to a solution of 3- hydroxybenzenesulfonamide (1.0 g, 5.8 mmol) and 3-(Boc-amino)propyl bromide (1.40 g, 5.8 mmol) in N,N-dimethylformamide (14 mL). The reaction was sealed with a yellow cap and heated at 55^°^C for 16 h. After cooling to room temperature, the reaction was diluted with water and isopropyl acetate. The aqueous layer was extracted with isopropyl acetate (3×30 mL). The combined organic layers were dried with sodium sulfate, concentrated and the crude residue was purified by flash column chromatography (silica, 100% isopropyl acetate) to give tert-butyl (3-(3-sulfamoylphenoxy)propyl)carbamate (1.34 g, 4.06 mmol, 70% Yield).

^1^H NMR (400 MHz, DMSO-*d*_6_) δ 7.47 (t, *J* = 7.9 Hz, 1H), 7.42–7.33 (m, 2H), 7.32 (s, 2H), 7.14 (ddd, *J* = 8.2, 2.5, 1.0 Hz, 1H), 6.90 (t, *J* = 5.6 Hz, 1H), 4.03 (t, *J* = 6.3 Hz, 2H), 3.09 (q, *J* = 6.6 Hz, 2H), 1.85 (p, *J* = 6.5 Hz, 2H), 1.37 (s, 9H).

^13^C NMR (101 MHz, DMSO) δ 159.07, 156.12, 145.84, 130.56, 118.48, 118.03, 111.84, 78.00, 66.17, 37.30, 29.54, 28.72.

HRMS (ESI) calcd for C_14_H_22_N_2_O_5_S [M-H]^+^: 329.1177, found: 329.1179.

#### Preparation of 4-(4-(Prop-2-yn-1-yloxy)benzoyl)-*N*-(3-(3-sulfamoylphenoxy)propyl)benzamide

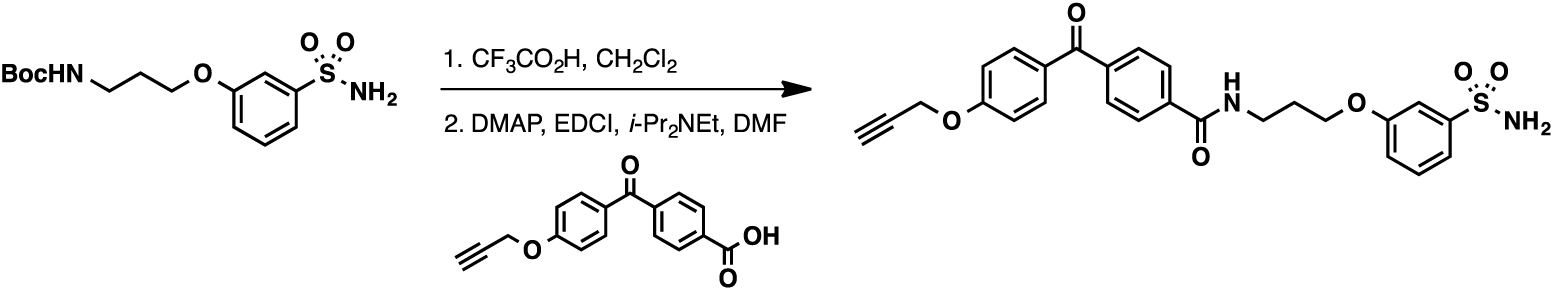

Trifluoroacetic acid (3.40 mL, 45.4 mmol) was added to a solution of tert-butyl (3-(3-sulfamoylphenoxy)propyl)carbamate (0.750 g, 2.27 mmol) in dichloromethane (45 mL) at room temperature. After 1 h, the reaction was concentrated under reduced pressure. Toluene (50 mL) was added and the reaction was again concentrated under reduced pressure. The crude residue was then submitted to the next step without further purification.

4-Dimethylaminopyridine (6.8 mg, 0.054 mmol) was added to a solution of 3-(3- aminopropoxy)benzenesulfonamide (92 mg, 0.401 mmol), 4-(4-prop-2-ynoxybenzoyl)benzoic acid (75 mg, 0.268 mmol), 1-(3-dimethylaminopropyl)-3-ethylcarbodiimide hydrochloride (62.2 mg, 0.321 mmol) and N,N-diisopropylethylamine (0.140 mL, 0.803 mmol) in N,N- dimethylformamide (2.7 mL) at room temperature. After 3 days, the reaction was diluted with water and isopropyl acetate and the aqueous layer was acidified to pH 1 using 1M HCl. The aqueous layer was extracted with isopropyl acetate (3×50 mL). The combined organic layers were dried with sodium sulfate, concentrated and the crude residue was purified by flash column chromatography (silica, 100% isopropyl acetate) to give 4-(4-(prop-2-yn-1-yloxy)benzoyl)-*N*-(3- (3-sulfamoylphenoxy)propyl)benzamide (46 mg, 0.093 mmol, 35% Yield).

^1^H NMR (400 MHz, DMSO-*d*_6_) δ 8.76 (t, *J* = 5.6 Hz, 1H), 8.03–7.96 (m, 2H), 7.82–7.73 (m, 4H), 7.48 (t, *J* = 7.9 Hz, 1H), 7.43–7.35 (m, 2H), 7.34 (s, 2H), 7.21–7.12 (m, 3H), 4.94 (d, *J* = 2.4 Hz, 2H), 4.12 (t, *J* = 6.1 Hz, 2H), 3.65 (t, *J* = 2.3 Hz, 1H), 3.49 (q, *J* = 6.4 Hz, 2H), 2.04 (p, *J* = 6.5 Hz, 2H).

^13^C NMR (101 MHz, DMSO) δ 194.49, 166.12, 161.52, 159.08, 145.86, 140.25, 137.95, 132.59, 130.60, 130.26, 129.62, 127.76, 118.51, 118.07, 115.29, 111.88, 79.27, 79.13, 66.29, 56.26, 36.83, 29.21.

HRMS (ESI) calcd for C_26_H_25_N_2_O_6_S [M+H]^+^: 493.1428, found: 493.1418.

#### Preparation of 4-Isocyanato-1,2,3,5,6,7-hexahydro-s-indacene

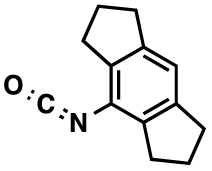

Triphosgene (88 mg, 0.288 mmol) was added in one portion to a solution of 1,2,3,5,6,7- hexahydro-s-indacen-4-amine (0.150 g, 0.866 mmol) and triethylamine (0.130 mL, 0.91 mmol) in THF (2.90 mL) and the mixture was heated to reflux. After 2 h the reaction was cooled to room temperature and the volatiles were removed under reduced pressure. The crude residue was dissolved in pentane and filtered through a plug of silica gel (to remove the triethylammonium chloride). The filtrate was concentrated under reduced pressure and the crude residue was submitted to the next step without further purification.

#### Preparation of Sodium ((1,2,3,5,6,7-hexahydro-*s*-indacen-4-yl)carbamoyl)((3-(3-(4-(4-(prop-2- yn-1-yloxy)-benzoyl)benzamido)propoxy)phenyl)sulfonyl)amide

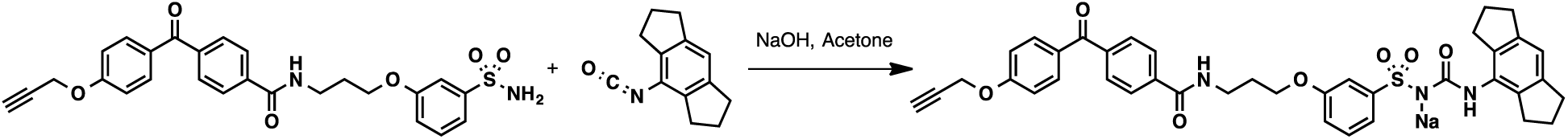

Sodium hydroxide (10 wt% in water, 0.040 mL, 0.102 mmol) was added to a solution of 4-(4- (prop-2-yn-1-yloxy)benzoyl)-*N*-(3-(3-sulfamoylphenoxy)propyl)benzamide (50 mg, 0.102 mmol) in acetone and the reaction was heated at reflux for 20 min. The reaction was cooled to room temperature and the volatiles were removed under reduced pressure. The crude residue was then placed on a high vacuum for 1 h to remove residual water, then it was dissolved in acetone (1.0 mL) and the reaction was heated at reflux. A solution of 4-isocyanato-1,2,3,5,6,7-hexahydro-s-indacene (27 mg, 0.137 mmol) in acetone (0.4 mL total with rinses) was added dropwise to the refluxing solution over 5 min. After addition, the reaction was allowed to reflux for an additional 10 min. The reaction was cooled to room temperature and the solvent was removed via syringe. Acetone (2.0 mL) was added to the solid, the mixture was sonicated for 10 sec and the solvent was removed via syringe. This sequence was repeated twice using heptane. The solid was isolated and dried under vacuum to give sodium ((1,2,3,5,6,7-hexahydro-*s*-indacen-4-yl)carbamoyl)((3-(3- (4-(4-(prop-2-yn-1-yloxy)benzoyl)benzamido)propoxy)phenyl)sulfonyl)amide: (32 mg, 0.045 mmol, 44% Yield).

^1^H NMR (400 MHz, DMSO-*d*_6_) δ 8.78 (t, *J* = 5.5 Hz, 1H), 8.03–7.95 (m, 2H), 7.81–7.70 (m, 4H), 7.41–7.35 (m, 2H), 7.32 (dt, *J* = 7.7, 1.4 Hz, 1H), 7.25 (t, *J* = 7.9 Hz, 1H), 7.19–7.10 (m, 2H), 6.92 (ddd, *J* = 8.0, 2.6, 1.2 Hz, 1H), 6.73 (s, 1H), 4.93 (d, *J* = 2.4 Hz, 2H), 4.06 (t, *J* = 6.2 Hz, 2H), 3.64 (t, *J* = 2.4 Hz, 1H), 3.46 (q, *J* = 6.5 Hz, 2H), 2.71 (t, *J* = 7.4 Hz, 4H), 2.63 (t, *J* = 7.3 Hz, 4H), 2.02 (q, *J* = 6.4 Hz, 2H), 1.87 (p, *J* = 7.5 Hz, 4H).

^13^C NMR (101 MHz, DMSO) δ 194.50, 166.10, 161.51, 159.18, 158.42, 149.29, 142.59, 140.19, 137.94, 137.28, 133.01, 132.59, 130.27, 129.59, 129.18, 127.79, 118.76, 116.20, 116.14, 115.28, 112.55, 79.27, 79.13, 65.92, 56.27, 36.91, 33.05, 30.98, 29.33, 25.53.

HRMS (ESI) calcd for C_39_H_38_N_3_O_7_S [M+H]^+^: 692.2415, found: 692.2425.

### Synthesis of iBodies

The HPMA polymer conjugates, named iBody U-121 and U-126, were prepared by reaction of the polymer precursor poly(HPMA-co-Ma-ß-Ala-TT) with a combination of the affinity anchor *N*-(2- aminoethyl)biotinamide hydrobromide (biotin-NH2) and the targeting ligand. Polymer precursors poly(HPMA-co-Ma-ß-Ala-TT) were synthesized as described earlier (Šubr and Ulbrich, 2006). In iBody U-121, the targeting ligand is compound 2, in which MCC950 is connected to a linker, compound 1, prepared as described (Simon et al., 2018). In iBody U-126, the targeting ligand is the GCPII inhibitor modified with a diazirine linker, which is prepared as described (Simon et al., 2018).

Preparation of compound 2. MCC950 (sodium salt, 52 mg, 0.122 mmol) and compound 1 (62 mg, 0.066 mmol) were dissolved in methanol and evaporated to dryness *in vacuo*. The solid residue was irradiated using LED diode (365 nm, 1 W) until starting compound 1 was present (UPLC/MS analysis). The reaction mixture was dissolved in 80% methanol and subjected to Sephadex LH-20 gel chromatography. Fractions containing desired product (UPLC/MS analysis) were collected, evaporated to dryness, and purified using preparative HPLC. Fractions containing desired product (UPLC/MS analysis) were collected, evaporated to dryness, and used in reaction with HPMA polymer. Mixture of isomers compound 2 was isolated as white solid (5 mg). UPLC/MS (m/z) 655.33 [M+2H]^++^, Tr 3.56-4.19 min; HRMS (ESI): Calculated for C_61_H_96_F_3_N_4_O_21_S [M+H]^+^: 1309.6240, Found: 1309.6220 (Supplemental Figure S2).

Preparation of iBody U-121. Polymer precursor poly(HPMA-*co*-Ma-β-Ala-TT) (15 mg), compound 2 (PSI137g3) (3.0 mg) and biotin-NH_2_ (1.5 mg) were dissolved in 0.3 ml DMSO and *N*,*N*-diisopropylethylamine (DIPEA) (2.3 µl) was added. The reaction mixture was stirred at room temperature for 4.5 h and then 2 µl of 1-aminopropan-2-ol was added and the reaction was again stirred for 10 min. Solution of polymer conjugate poly(HPMA-*co*-Ma-β-Ala-compound 2-Ma-β- Ala-NH-biotin) was diluted with 0.7 ml of methanol and purified on chromatography column Sephadex LH-20 in methanol equipped with UV detector (220 nm). Polymer conjugate was dissolved in 1.5 ml of distilled water and lyophilized. Yield of U-121 (M_n_ = 303,000 g/mol, M_w_ = 982,000 g/mol, Ð = 3.24) was 13 mg, content of biotin was 3.3 wt% (Supplemental Figure S2).

Preparation of iBody U-126. Polymer precursor poly(HPMA-*co*-Ma-β-Ala-TT) (10 mg), GCPII inhibitor modified with a diazirine linker (3.0 mg), biotin-NH_2_ (1.1 mg) and ATTO488-NH_2_ (0.8 mg) were dissolved in 0.5 ml DMSO, followed by the addition of DIPEA (5.2 µl). Reaction was carried out for 4.5 h at room temperature and then 2 µl of 1-aminopropan-2-ol was added and the reaction was stirred for 10 min. Solution of polymer conjugate was diluted with 1.5 ml of methanol and purified on chromatography column Sephadex LH-20 in methanol. Methanol was evaporated and polymer conjugate was dissolved in 1.5 ml of distilled water and purified on chromatography column Sephadex G-25 and lyophilized. Yield of U-126 (M_n_ = 193,500 g/mol, M_w_ = 685,000 g/mol, Ð = 3.54) was 9 mg, content of biotin was 3.5 wt% (Supplemental Figure 3).

## Supplementary Figure legends

**Figure S1.**
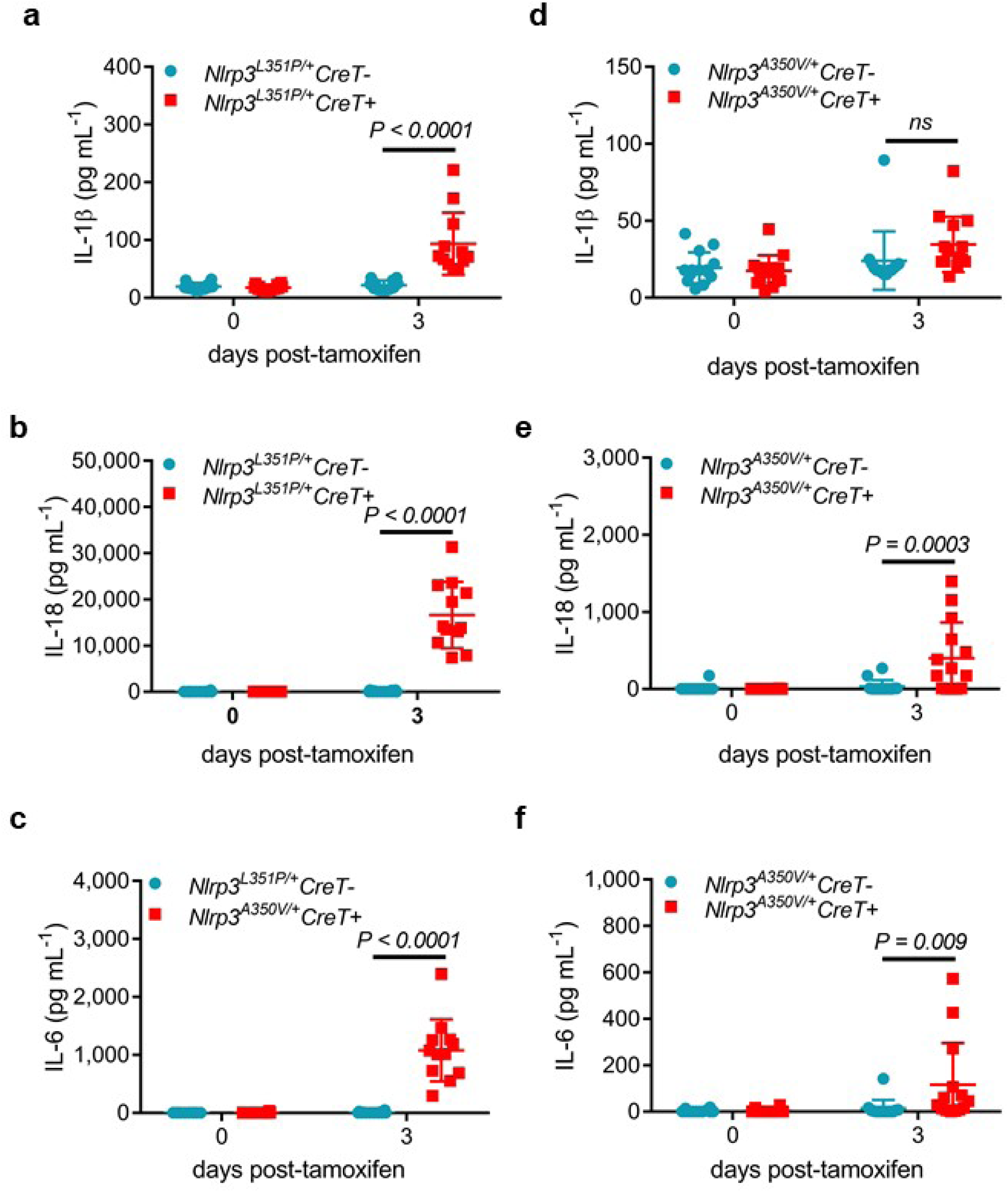
Tamoxifen-inducible mouse models of CAPS. (A-C) Serum levels of IL-1β (A), IL-18 (B) and IL-6 (C) in *Nlrp3^L351P/+^ CreT+* and *CreT-* littermates before and after tamoxifen treatment at day 3. (D-E) Serum levels of IL-1β (D), IL-18 (E) and IL-6 (F) in *Nlrp3^A350V/+^CreT+* and *CreT-* littermates before and after tamoxifen treatment at day 3.

**Figure S2.**
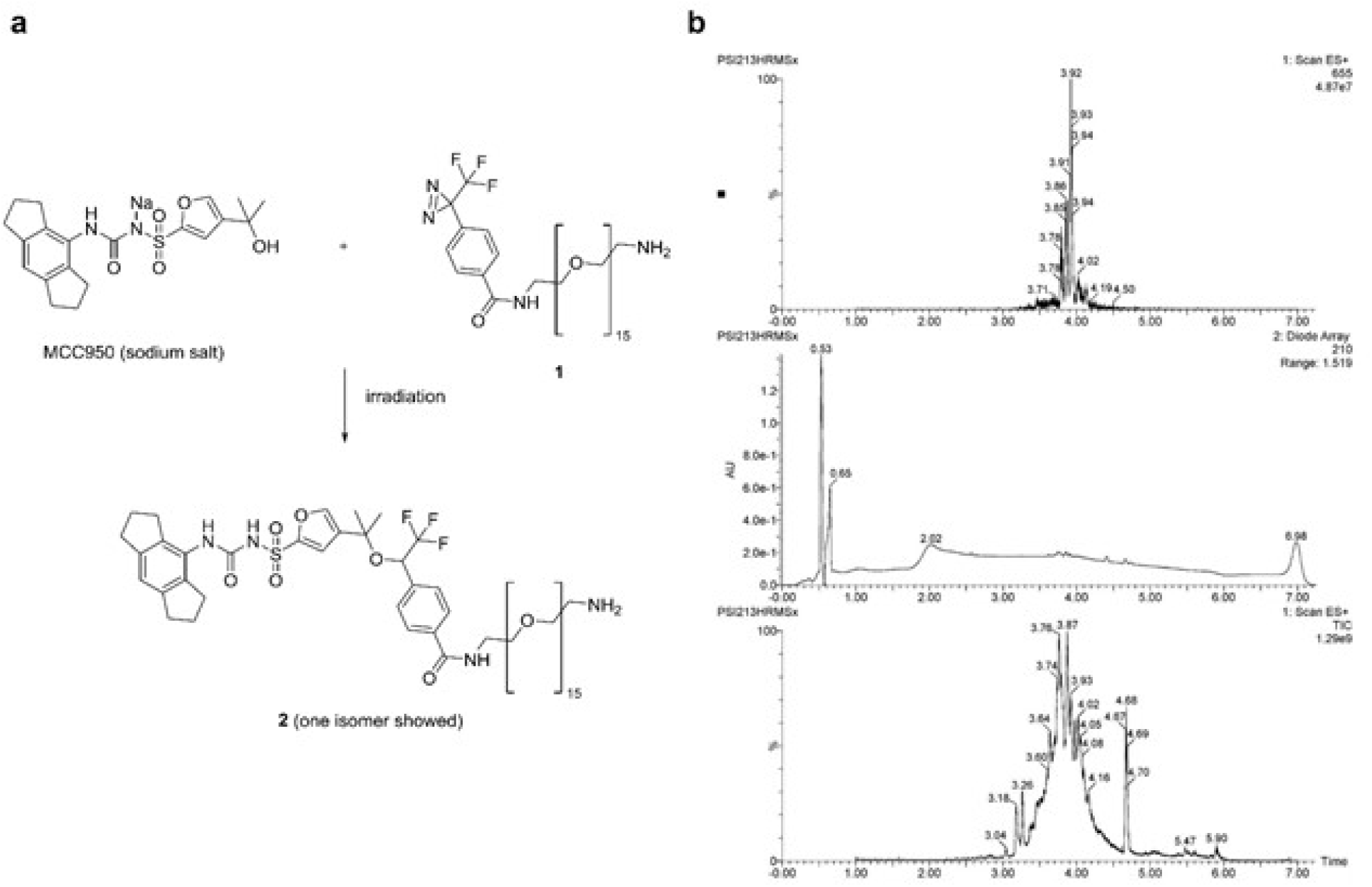
Synthesis of compound 2. (A-B) Synthesis (A) and UPLC/MS analysis (B) of mixture of isomers compound 2 (PSI213) (total ion chromatogram, 210 nm chromatogram, mass [655]^++^ chromatogram).

**Figure S3.**
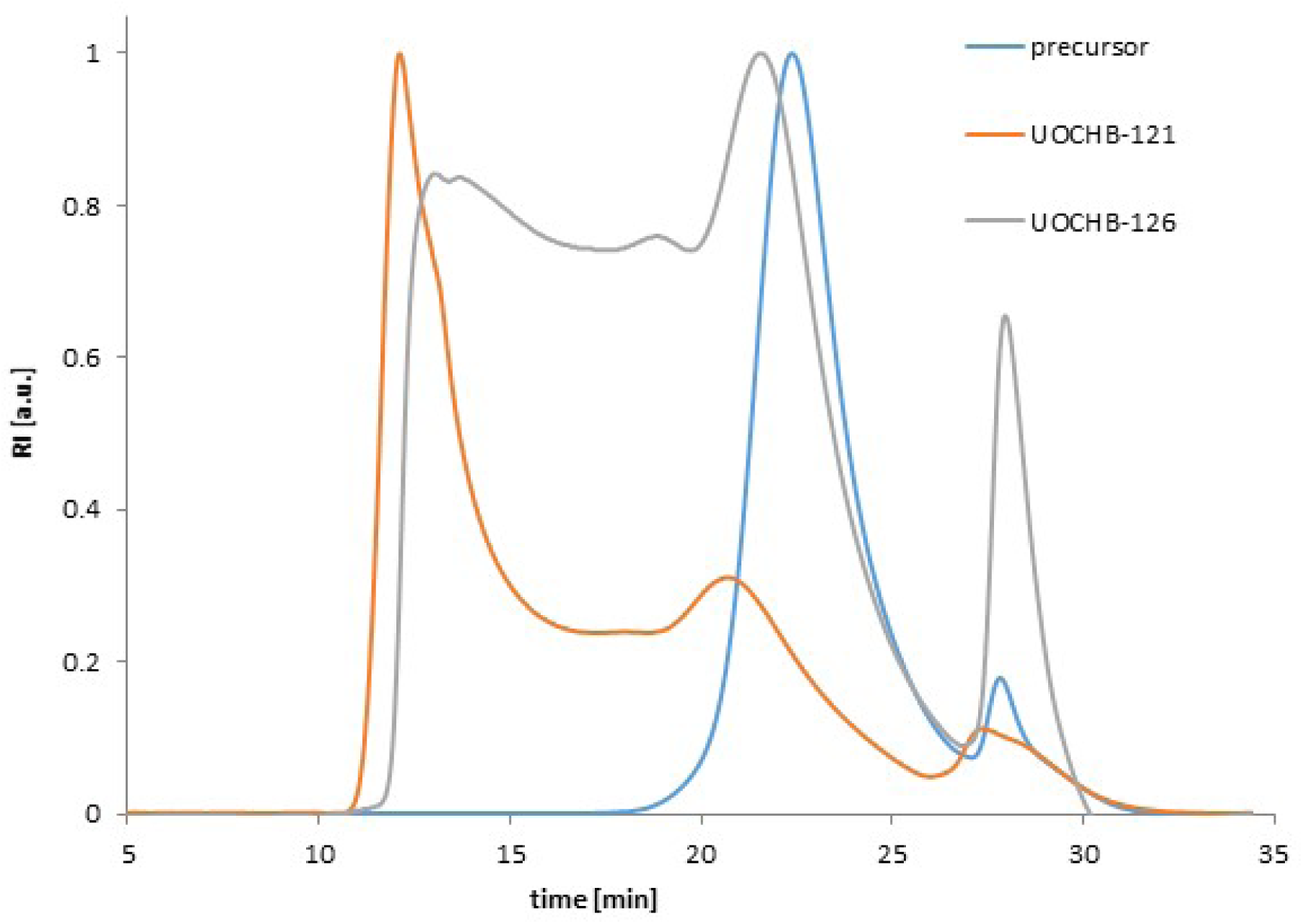
GPC chromatogram of polymer precursor (blue), polymer conjugate U-121 (orange) and U-126 (grey).

## References

1. Bauernfeind, F.G., Horvath, G., Stutz, A., Alnemri, E.S., MacDonald, K., Speert, D., Fernandes-Alnemri, T., Wu, J., Monks, B.G., Fitzgerald, K.A., et al. (2009). Cutting edge: NF-kappaB activating pattern recognition and cytokine receptors license NLRP3 inflammasome activation by regulating NLRP3 expression. Journal of immunology 183, 787–791.

2. Broz, P., and Dixit, V.M. (2016). Inflammasomes: mechanism of assembly, regulation and signalling. Nat Rev Immunol 16, 407–420.

3. Brydges, S.D., Mueller, J.L., McGeough, M.D., Pena, C.A., Misaghi, A., Gandhi, C., Putnam, C.D., Boyle, D.L., Firestein, G.S., Horner, A.A., et al. (2009). Inflammasome-mediated disease animal models reveal roles for innate but not adaptive immunity. Immunity 30, 875–887.

4. Coll, R.C., Robertson, A.A., Chae, J.J., Higgins, S.C., Munoz-Planillo, R., Inserra, M.C., Vetter, I., Dungan, L.S., Monks, B.G., Stutz, A., et al. (2015). A small-molecule inhibitor of the NLRP3 inflammasome for the treatment of inflammatory diseases. Nature medicine 21, 248–255.

5. Duncan, J.A., Bergstralh, D.T., Wang, Y., Willingham, S.B., Ye, Z., Zimmermann, A.G., and Ting, J.P. (2007). Cryopyrin/NALP3 binds ATP/dATP, is an ATPase, and requires ATP binding to mediate inflammatory signaling. Proc Natl Acad Sci U S A 104, 8041–8046.

6. Grinstein, L., Endter, K., Hedrich, C.M., Reinke, S., Luksch, H., Schulze, F., Robertson, A.A.B., Cooper, M.A., Rosen-Wolff, A., and Winkler, S. (2018). An optimized whole blood assay measuring expression and activity of NLRP3, NLRC4 and AIM2 inflammasomes. Clin Immunol 191, 100–109.

7. Hagar, J.A., Powell, D.A., Aachoui, Y., Ernst, R.K., and Miao, E.A. (2013). Cytoplasmic LPS activates caspase-11: implications in TLR4-independent endotoxic shock. Science 341, 1250–1253.

8. He, Y., Franchi, L., and Nunez, G. (2013). TLR agonists stimulate Nlrp3-dependent IL-1beta production independently of the purinergic P2X7 receptor in dendritic cells and in vivo. Journal of immunology 190, 334–339.

9. Kayagaki, N., Stowe, I.B., Lee, B.L., O’Rourke, K., Anderson, K., Warming, S., Cuellar, T., Haley, B., Roose-Girma, M., Phung, Q.T., et al. (2015). Caspase-11 cleaves gasdermin D for non-canonical inflammasome signalling. Nature 526, 666–671.

10. Kayagaki, N., Warming, S., Lamkanfi, M., Vande Walle, L., Louie, S., Dong, J., Newton, K., Qu, Y., Liu, J., Heldens, S., et al. (2011). Non-canonical inflammasome activation targets caspase-11. Nature 479, 117–121.

11. Kayagaki, N., Wong, M.T., Stowe, I.B., Ramani, S.R., Gonzalez, L.C., Akashi-Takamura, S., Miyake, K., Zhang, J., Lee, W.P., Muszynski, A., et al. (2013). Noncanonical inflammasome activation by intracellular LPS independent of TLR4. Science 341, 1246–1249.

12. Laliberte, R.E., Perregaux, D.G., Hoth, L.R., Rosner, P.J., Jordan, C.K., Peese, K.M., Eggler, J.F., Dombroski, M.A., Geoghegan, K.F., and Gabel, C.A. (2003). Glutathione s-transferase omega 1- 1 is a target of cytokine release inhibitory drugs and may be responsible for their effect on interleukin-1beta posttranslational processing. J Biol Chem 278, 16567–16578.

13. Lamkanfi, M., and Dixit, V.M. (2014). Mechanisms and functions of inflammasomes. Cell 157, 1013–1022.

14. Lamkanfi, M., Mueller, J.L., Vitari, A.C., Misaghi, S., Fedorova, A., Deshayes, K., Lee, W.P., Hoffman, H.M., and Dixit, V.M. (2009). Glyburide inhibits the Cryopyrin/Nalp3 inflammasome. J Cell Biol 187, 61–70.

15. Mackinnon, A.L., and Taunton, J. (2009). Target Identification by Diazirine Photo-Cross-linking and Click Chemistry. Curr Protoc Chem Biol 1, 55–73.

16. Mangan, M.S.J., Olhava, E.J., Roush, W.R., Seidel, H.M., Glick, G.D., and Latz, E. (2018). Targeting the NLRP3 inflammasome in inflammatory diseases. Nat Rev Drug Discov 17, 688.

17. Mariathasan, S., Weiss, D.S., Newton, K., McBride, J., O’Rourke, K., Roose-Girma, M., Lee, W.P., Weinrauch, Y., Monack, D.M., and Dixit, V.M. (2006). Cryopyrin activates the inflammasome in response to toxins and ATP. Nature 440, 228–232.

18. Matusiak, M., Van Opdenbosch, N., Vande Walle, L., Sirard, J.C., Kanneganti, T.D., and Lamkanfi, M. (2015). Flagellin-induced NLRC4 phosphorylation primes the inflammasome for activation by NAIP5. Proc Natl Acad Sci U S A 112, 1541–1546.

19. McGeough, M.D., Pena, C.A., Mueller, J.L., Pociask, D.A., Broderick, L., Hoffman, H.M., and Brydges, S.D. (2012). Cutting edge: IL-6 is a marker of inflammation with no direct role in inflammasome-mediated mouse models. Journal of immunology 189, 2707–2711.

20. Munoz-Planillo, R., Kuffa, P., Martinez-Colon, G., Smith, B.L., Rajendiran, T.M., and Nunez, G. (2013). K(+) efflux is the common trigger of NLRP3 inflammasome activation by bacterial toxins and particulate matter. Immunity 38, 1142–1153.

21. Primiano, M.J., Lefker, B.A., Bowman, M.R., Bree, A.G., Hubeau, C., Bonin, P.D., Mangan, M., Dower, K., Monks, B.G., Cushing, L., et al. (2016). Efficacy and Pharmacology of the NLRP3 Inflammasome Inhibitor CP-456,773 (CRID3) in Murine Models of Dermal and Pulmonary Inflammation. Journal of immunology 197, 2421–2433.

22. Qu, Y., Misaghi, S., Izrael-Tomasevic, A., Newton, K., Gilmour, L.L., Lamkanfi, M., Louie, S., Kayagaki, N., Liu, J., Komuves, L., et al. (2012). Phosphorylation of NLRC4 is critical for inflammasome activation. Nature 490, 539–542.

23. Ridker, P.M., Everett, B.M., Thuren, T., MacFadyen, J.G., Chang, W.H., Ballantyne, C., Fonseca, F., Nicolau, J., Koenig, W., Anker, S.D., et al. (2017). Antiinflammatory Therapy with Canakinumab for Atherosclerotic Disease. N Engl J Med 377, 1119–1131.

24. Shi, J., Zhao, Y., Wang, K., Shi, X., Wang, Y., Huang, H., Zhuang, Y., Cai, T., Wang, F., and Shao, F. (2015). Cleavage of GSDMD by inflammatory caspases determines pyroptotic cell death. Nature 526, 660–665.

25. Shi, J., Zhao, Y., Wang, Y., Gao, W., Ding, J., Li, P., Hu, L., and Shao, F. (2014). Inflammatory caspases are innate immune receptors for intracellular LPS. Nature 514, 187–192.

26. Simon, P., Knedlik, T., Blazkova, K., Dvorakova, P., Brezinova, A., Kostka, L., Subr, V., Konvalinka, J., and Sacha, P. (2018). Identification of Protein Targets of Bioactive Small Molecules Using Randomly Photomodified Probes. ACS Chem Biol 13, 3333–3342.

27. Smith, E., and Collins, I. (2015). Photoaffinity labeling in target- and binding-site identification. Future Med Chem 7, 159–183.

28. Van Gorp, H., Van Opdenbosch, N., and Lamkanfi, M. (2019). Inflammasome-Dependent Cytokines at the Crossroads of Health and Autoinflammatory Disease. Cold Spring Harb Perspect Biol 11.

29. Ventura, A., Kirsch, D.G., McLaughlin, M.E., Tuveson, D.A., Grimm, J., Lintault, L., Newman, J., Reczek, E.E., Weissleder, R., and Jacks, T. (2007). Restoration of p53 function leads to tumour regression in vivo. Nature 445, 661–665.

30. Voet, S., Srinivasan, S., Lamkanfi, M., and van Loo, G. (2019). Inflammasomes in neuroinflammatory and neurodegenerative diseases. EMBO Mol Med.

31. Wang, G.G., Calvo, K.R., Pasillas, M.P., Sykes, D.B., Hacker, H., and Kamps, M.P. (2006). Quantitative production of macrophages or neutrophils ex vivo using conditional Hoxb8. Nat Methods 3, 287–293.

## References to supplementary data

Simon, P., Knedlik, T., Blazkova, K., Dvorakova, P., Brezinova, A., Kostka, L., Subr, V., Konvalinka, J., and Sacha, P. (2018). Identification of Protein Targets of Bioactive Small Molecules Using Randomly Photomodified Probes. ACS Chem Biol 13, 3333–3342.

Šubr, V., and Ulbrich, K. (2006). Synthesis and properties of new N-(2-hydroxypropyl) methacrylamide copolymers containing thiazolidine-2-thione reactive groups. Reactive and Functional Polymers 66, 1525–1538.

